# Does ecological restoration align prokaryotic community structure with natural references across European coastal wetlands?

**DOI:** 10.64898/2025.12.26.696618

**Authors:** A Picazo, C Rochera, D Morant, M Pedrón, M Adamescu, R Ambrosio, K Attermeyer, C Santinelli, C Tropea, Gf Bachi, N Bègue, M Bučas, M Cabrera-Brufau, A Camacho Santamans, R Carballeira, M Carloni, L Cavalcante, C Cazacu, G Checcucci, J.P Coelho, V Dinu, V Evangelista, J Gintauskas, R Giuca, A Guelmami, M Guerrazzi, S Hilaire, M Kataržytė, A.I Lillebø, R Lizán, C Minaudo, B Misteli, J Montes-Pérez, B Obrador, B.R.F Oliveira, V.H Oliveira, J Petkuvienė, T Racoviceanu, S Ribeiro, M Ronse, A Sousa, W Suykerbuyk, E Tiškus, D Vaičiūtė, S Valsecchi, M van Puijenbroek, O Andreu, M Yegon, D von Schiller, A Camacho

**Affiliations:** Institut Cavanilles de Biodiversitat i Biologia Evolutiva, Universitat de València, C/Catedràtic José Beltran 2, 46980, Paterna, Spain; Departament de Biologia Evolutiva, Ecologia i Ciències Ambientals, Universitat de Barcelona, Av. Diagonal 643, 8028, Barcelona, Spain; WasserCluster Lunz - Biologische Station, University of Vienna, Dr. Carl Kupelwieser Promenade 5, 3293, Lunz am See, Austria; Department of Systems Ecology and Sustainability, University of Bucharest, spl. Independentei 91 - 95, 50095, Bucharest, Romania; Institut de recherche pour la conservation des zones humides méditerranéennes, Tour du Valat, Le Sambuc, 13200, Arles, France; Istituto di Biofisica, Consiglio Nazionale delle Ricerche (CNR) Pisa, Via Giuseppe Moruzzi 1, 56124, Pisa, Italy; Marine Research Institute, Klaipėda University, Universiteto ave. 17, 92294, Klaipėda, Lithuania; ECOMARE, CESAM - Centre for Environmental and Marine Studies, Department of Biology, University of Aveiro, Campus de Santiago, 3810-193, Aveiro, Portugal; Wageningen Marine Research, Wageningen University & Research, Korringaweg 7, 4401 NT, Yerseke, The Netherlands; Departamento de Biología Vegetal, Facultad de Farmacia, Universitat de València, 46100 Burjassot, Spain

**Keywords:** Next-generation sequencing, 16S rRNA metabarcoding, Microbial community composition, Indicator species, European Coastal Wetlands

## Abstract

Coastal wetlands are crucial for biodiversity and act as critical buffers for carbon sequestration and atmospheric greenhouse gases (GHG) concentrations, yet their degradation often turns them into GHG sources. Restoration is widely implemented to recover these services, but it remains unclear whether interventions successfully reestablish the microbial functional diversity underpinning biogeochemical cycles. We tested the hypothesis that restoration aligns prokaryotic community structure with natural references, analyzing, a European gradient of coastal wetlands, comparing well-preserved, altered, and restored sites in water and sediment. Using 16SrRNA-metabarcoding and IndVal-Analysis, we characterized community assembly identifying diagnostic functional consortia. Results revealed a marked difference in water and sediment recovery after restoration. Bacterioplankton communities rapidly converge to natural references, while sediment microbiome displayed significant “ecological memory”. Restored wetlands show sediment communities structurally distinct from well-preserved, retaining alteration-associated guilds decades. Results support the initial hypothesis: restoration processes in coastal wetlands can re-establish communities and metabolisms resembling well-preserved conditions in the water in the short term, while sediments retain microbial communities and metabolisms inherited from altered conditions for a long time. Future strategies must integrate active sediment interventions using molecular bioindicators to validate not only the landscape appearance, but the effective reactivation of ecosystem processes and microbiota-related services.

## Introduction

Coastal wetlands represent critical ecotones at the land-ocean interface, playing a disproportionately high role in regulating global biogeochemical cycles despite their limited geographic extent (Airoldi & Beck, 2007; Davidson et al., 2018). These ecosystems, ranging from marshes and lagoons to deltas and estuaries, are renowned for their exceptional capacity to sequester and store organic carbon, acting as key providers of regulating ecosystem services (Mitsch et al., 2015; IUCN, 2021). However, their strategic position and high productivity also render them extremely vulnerable to anthropogenic pressures and global environmental change (Newton et al., 2020). The alteration or degradation of these systems can shift their function from net carbon sinks to significant sources of greenhouse gases (GHG), specifically carbon dioxide (CO_2_), methane (CH_4_), and nitrous oxide (N_2_O) (Mitsch et al., 2014; Camacho et al., 2017; Campbell et al., 2025;). In this context, ecological restoration projects have emerged as a priority strategy, not only to recover lost biodiversity but fundamentally to re-establish the biogeochemical functionality that ensures climate resilience and the reversal of carbon fluxes towards mitigation (Moreno-Mateos et al., 2015; Kikstra et al., 2022; Misteli et al. under review).

To understand the dynamics of the processes associated with the biogeochemical functionality of the ecosystem on the European continent, it is imperative to categorize the vast heterogeneity of these wetlands into functional units that allow for robust comparative analysis. In this context, three major hydrological and ecological typologies dominating the European coastal landscape can be distinguished: firstly, Mediterranean coastal marshes and wetlands, characterized by often temporary hydrology and intensive artificial water management (Morant et al., 2020); secondly, Atlantic intertidal ecosystems, strongly influenced by tidal dynamics and salinity gradients; and finally, large lagoon or deltaic systems of permanent waters, which act as transition and buffer zones between major river basins and the marine environment (Vybernaite-Lubiene et al., 2022). The variability in hydrological regimes and salinity among different wetlands groups determine not only the structure of halophytic and/or submerged vegetation but fundamentally the underlying biogeochemical processes (Camacho-Santamans et al., 2024). Given that these transformations are largely orchestrated by the metabolic machinery of prokaryotic communities, an in-depth study of microbial diversity proves indispensable for deciphering the biotic mechanisms that regulate the main biogeochemical cycles and ultimately GHG emissions (Neubauer & Megonigal, 2021; Cao et al., 2024).

In response to the historical and ongoing degradation of these ecosystems, ecological restoration has emerged as a priority strategy in global and European environmental policy (De Stefano et al., 2023), aligning with key initiatives such as the UN Decade on Ecosystem Restoration and the Regulation (EU) 2024/1991 on nature restoration. Restoration approaches are multifaceted, ranging from passive hydrological restoration and tidal reconnection to active interventions involving revegetation, morphological re-profiling, or the remediation of eutrophication and pollution (Gattuso et al., 2020). The value of these efforts extends far beyond the mere recovery of vegetation cover or landscape aesthetics; their true success must be gauged by the restitution of ecosystem functionality. However, restoration interventions inevitably induce significant physicochemical disturbances that shift redox conditions, nutrient availability, and substrate structure. These modifications exert a direct and immediate impact on the ecosystem’s most dynamic functional component: the microbial community (Allison & Martiny, 2008). Since microorganisms respond to environmental shifts much faster than plants or macrofauna, changes in wetland management have the potential to rapidly reconfigure microbial food webs, thereby altering the metabolic pathways that govern the fate of carbon and nutrients in both aquatic and sedimentary compartments (Prosser et al., 2007).

Microbial communities are fundamental to global ecosystem functioning, particularly regarding wetland carbon, GHG fluxes and C mass balance, as they act as the catalytic engines of biogeochemical cycles. CH4 emission, for instance, is the net result of the balance between its production by methanogenic archaea under strict anaerobic conditions and its consumption by methanotrophic bacteria at oxic-anoxic interfaces (Canfield et al., 2005, Morant et al., 2024). Similarly, N_2_O emissions are regulated by complex microbial consortia mediating respiration, denitrification, and nitrification processes (Ussiri & Lal, 2013). Therefore, a detailed understanding of microbial diversity is essential to elucidate the mechanisms driving GHG fluxes at the ecosystem level. Historically, methodological constraints prevented a deep characterization of these communities; however, the advent and standardization of high-throughput molecular techniques, such as 16S rRNA gene amplicon sequencing (metabarcoding), have revolutionized our capacity to estimate alpha and beta diversity, as well as the taxonomic structure of prokaryotic communities (Quast et al., 2012; Borja et al., 2019). The application of these molecular tools allows not only for the inventorying of the immense biodiversity hidden in wetland water and sediments (Thompson et al., 2017) but also for the inference of potential metabolic functions and the understanding of how environmental disturbances, including those associated with restoration, shape the architecture of these invisible yet functionally critical communities (Bier et al., 2015).

In the context of restoration assessment, the identification of specific microbial genera or microbial functional guilds acting as bioindicators is of critical strategic importance. Beyond merely describing general shifts in diversity, the inclusion of microbial indicators leads to a more comprehensive wetland assessment for restoration and management (Sims et al., 2013), allowing for the diagnosis of the ecosystem’s ‘metabolic’ state by revealing invisible processes, such as active methanogenesis or incipient denitrification, before they become evident in water chemistry or vegetation. Establishing significant taxonomic and functional differences between well-preserved (reference), altered (degraded), and restored sites is crucial for validating the effectiveness of interventions (Dufrêne & Legendre, 1997). The central hypothesis of this comparative approach posits that if restoration is effective, the microbial community structure of the restored site should diverge from the dysfunctional configuration of the altered site and converge towards that of the well-preserved site, thereby recovering essential ecosystem functions (Campbell et al., 2025). However, the resilience of altered communities and the potential existence of alternative stable states may generate incomplete or hybrid recovery trajectories. Therefore, the comparative analysis of microbial diversity and composition across these three conservation states (well-preserved, altered, restored) throughout different European coastal wetland typologies offers a unique opportunity. This approach not only facilitates an understanding of post-restoration community assembly rules but, by identifying which specific species lead the recovery (or resistance to change), enables the development of precise molecular monitoring tools to audit the true long-term functional success of restoration.

The present study addresses restoration effect on European coastal wetlands through a comprehensive analysis of microbial diversity across multiple pilot sites in Europe. The central hypothesis posits that ecological restoration reconfigures the diversity and complexity of the microbial community, in both water and sediment, steering it towards states structurally and functionally analogous to natural references. To test this hypothesis, the following specific objectives are proposed: (1) to assess the impact of restoration on prokaryotic community composition in water and sediments by contrasting them with those of altered and well-preserved sites; (2) to identify indicator prokaryotic indicator groups that diagnose the success or stagnation of restoration trajectories; and (3) to evaluate the functional implications of these structural shifts, with special emphasis on metabolic pathways associated with GHG emission or mitigation. Through this integrated approach, this study aims to provide a mechanistic basis to optimize coastal wetland restoration strategies, ensuring that such interventions transcend the recovery of the visible landscape to re-establish essential biogeochemical functionality within the context of climate change.

## Material & Methods

### 2.1. Study area and experimental design

The study was conducted across six Case Pilots (CPs) distributed throughout Europe, selected to represent a broad gradient of coastal and transitional wetland ecosystems subject to diverse anthropogenic pressures and management strategies (Fig. 1). In the Mediterranean region, the Camargue (CA), (France), comprises a vast alluvial plain where a mosaic of natural wetlands (lagoons, marshes, steppes) coexists with intensive agro-ecosystems. This system is heavily regulated by a complex irrigation and drainage network that balances agricultural productivity with the conservation of Ramsar-protected habitats. Marjal dels Moros (VA), (Spain), constitutes a biodiversity reservoir restored in the 1990s within a historically pressured agricultural landscape. Protected under the Natura 2000 network, its management focuses on maintaining adequate flood levels (even if it means altering the natural drying cycle and therefore the seasonal patterns of salinity) to support endangered endemic flora and fauna, serving as a critical refuge in the Western Mediterranean.

**Figure 1.**
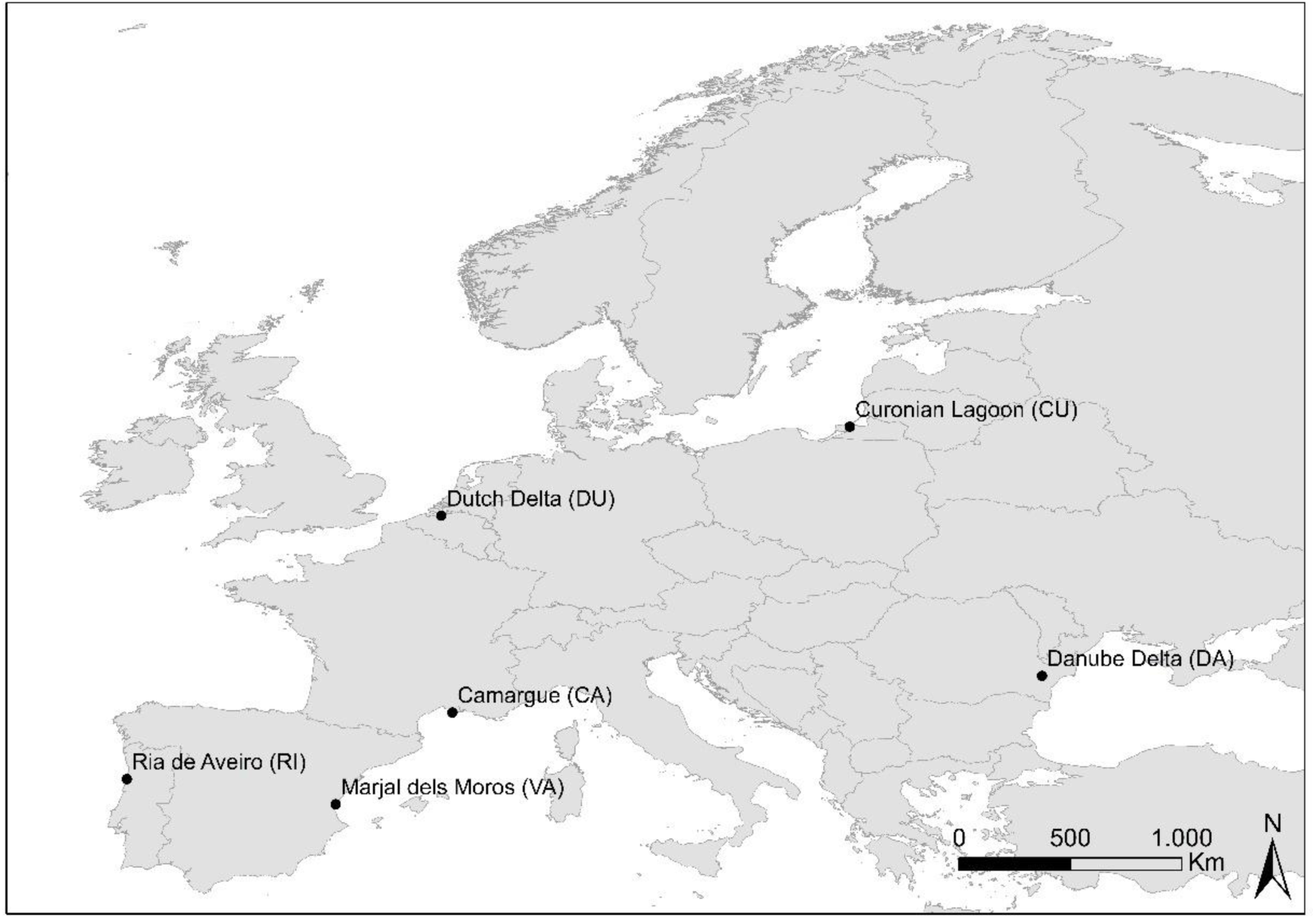
Sampling locations in the six European Case Pilots (Camargue (CA), Curonian Lagoon (CU), Ria de Aveiro (RI), Southwest Dutch Delta (DU), Danube Delta (DA), and Marjal del Moros (VA)).

On the Atlantic coast, the Ria de Aveiro (RI), (Portugal), encompasses intertidal *Zostera noltei* seagrass meadows, which provides key services, including blue carbon sequestration. However, this complex socio-ecological system faces significant pressures from industrial activity, dredging, and habitat fragmentation. In Northern Europe, the Southwest Dutch Delta (DU), (The Netherlands) exemplifies an estuarine system heavily modified by major hydraulic infrastructure at the confluence of the Rhine, Meuse, and Scheldt rivers. While it retains valuable intertidal flats and shoals, the region currently faces challenges related to “coastal squeeze,” erosion, and altered hydrodynamics. The Curonian Lagoon (CU), (Lithuania), Europe’s largest coastal lagoon, is a shallow, semi-enclosed system distinguished by the strong freshwater influence of the Nemunas River and the physical barrier of the Curonian Spit. It is highly productive but suffers from severe eutrophication and frequent cyanobacterial blooms, requiring management strategies that balance conservation with tourism and fisheries. Finally, the Danube Delta (DA), (Romania), represents one of the continent’s best-preserved deltas, hosting the world’s largest continuous reed bed expanses. It plays a pivotal role in the region’s biogeochemistry by acting as a natural filter for the Black Sea, regulating flood pulses, and sustaining exceptional biodiversity.

The experimental design was hierarchically structured to assess the impact of restoration at both continental and local scales (Table 1). Stratified design was implemented by selecting six subsites representing three conservation statuses: well-preserved (WP), altered (A), and restored (R), with two independent spatial replicates per status (e.g., WP1, WP2). To capture temporal dynamics, four seasonal sampling campaigns (S1 - autumn) to S4 -summer) were conducted. Notably, sampling effort was intensified during the spring campaign (S3), considered the period of ecological optimum and maximum biological activity; for this campaign, three biological replicates (n=3) were collected at each subsite and matrix (water and sediment) to capture intra-site variability, whereas the autumn (S1), winter (S2), and summer (S4) campaigns followed a standard monitoring strategy with a single replicate (n=1) per subsite and matrix (water and sediment).

**Table 1.** Overview of the case pilot sites. The table summarizes geographical location, ecosystem type, specific selected habitats, main anthropogenic pressures (alterations), and implemented restoration strategies.

| Case Pilot Site (Code) | Location / Region | Ecosystem Type | Selected Habitat for Sampling | Main Alteration Pressures | Restoration Approach |
| --- | --- | --- | --- | --- | --- |
| Camargue (CA) | France (Mediterranean) | Floodplain / Delta | Freshwater marshes and ponds | Hydrological change (rice cultivation) | Soil, hydrology, and morphological reconstruction |
| Curonian Lagoon (CU) | Lithuania (Baltic Sea) | Coastal Lagoon | Submerged plant beds (on sand/mud) | Eutrophication, mud accumulation | Passive recovery (reduced mud, increased vegetation) |
| Ria de Aveiro (RI) | Portugal (Atlantic) | Coastal Lagoon | Intertidal seagrass beds | Erosion, bait digging (bioturbation), pollution | Seagrass transplantation, physical protection |
| SW Dutch Delta (DU) | Netherlands (Atlantic) | Estuarine Delta | Intertidal salt marshes | Erosion, hydrodynamic reduction (breakwaters) | Managed realignment (active) & Dike failure (passive) |
| Danube Delta (DA) | Romania (Black Sea) | River Delta | Freshwater ponds with reed beds | Conversion to dryland (agriculture) | Hydrological reconnections |
| Marjal dels Moros (VA) | Spain (Mediterranean) | Coastal Wetlands | Brackish marshes | Hydrological, trophic, and morphological change | Soil, morphology, and vegetation recovery |

### 2.2. Environmental variables

To assess the physicochemical and trophic context of the ecosystems studied, key environmental variables were determined in the water and sediment following the standardized protocols of the RESTORE4Cs project (Oliveira et al., under review). In water, *in situ* parameters including water depth, temperature (Temp), dissolved oxygen (DO), and electrical conductivity (Cond) were measured. Additionally, alkalinity, bacterial biomass (TBacteria), photosynthetic biomass (Chlorophyll-*a*; Chl-*a*), and nutrients: orthophosphate (PO4), ammonium (NH4), and nitrate/nitrite (NOx) were analyzed, along with Total Nitrogen (Total-N) and Total Phosphorus (Total-P) content. In the sediment, characterization focused on the evaluation of soil moisture and ash-free dry mass, as well as the quantification of Total Organic Carbon percentage (TOC) and concentrations of Total Phosphorus (Total-P) and Total Nitrogen (Total-N). (Misteli et al., under review).

### 2.3. DNA extraction, 16S rRNA gene library preparation and sequencing

A total of 517 environmental samples were processed and sequenced, evenly distributed between water (n=222) and sediment (n=295) across the six European case pilots. For metabarcoding analysis of archaeal and bacterial communities, DNA extraction from water and sediment (300-500 mg) was performed with the EZNA Soil DNA isolation kit (Omega Bio-Tek, Inc., Norcross, GA, USA) following Picazo et al (2019). After quantification of each sample, sequencing of region V4 of the 16S rRNA gene was done using the Illumina MiSeq system (2×250bp). For each sample, Illumina compatible, dual indexed amplicon libraries of the 16S-V4 rRNA hypervariable region were created with primers 515f/806r. PCR reactions were made following Kozich et al., 2013. Completed libraries were normalized using Invitrogen SequalPrep DNA Normalization Plates. Then, the Qubit quantified pool was loaded on a standard Illumina MiSeq v2 flow cell and sequencing was performed in a 2×250 bp paired end format using a MiSeq v2 500 cycle reagent cartridge. Custom sequencing and index primers complementary to the 515/806 target sequences were added to appropriate wells of reagent cartridge. Base calling was done by Illumina Real Time Analysis (RTA) v1.18.54 and output of RTA was demultiplexed and converted to FastQ format with Illumina Bcl2fastq v2.19.1.

Sequences were processed using the UPARSE pipeline using USEARCH v12b (Edgar, 2013). After merging read pairs, the dataset was filtered by a maximum number of expected errors of 0.5%. Chimeric sequences were removed with UCHIME. Filtered sequences were clustered in zero-radius Operational Taxonomic Units (ZOTUs), which are sequences with 100% identity. The unoise3 method was performed for denoising sequencing errors in Illumina-sequenced amplicons (Edgar 2016). Alignment and taxonomic assignment were done with SINA v1.2.1152 using SILVA 138.1 database (Pruesse et al., 2012). SINA uses the Lowest Common Ancestor method (LCA). We configured a “Min identity” threshold of 0.8 and a maximum number of search results of 1 per sequence, resulting in “best match” type. Sequences with low alignment quality (<90%) and sequences identified as mitochondria or chloroplasts were removed from the analysis. Original ZOTU table were normalized by rarefying the reads of all samples, rarefactions were repeated 100 times to avoid the loss of less abundant ZOTUs, and the rarefactions were unified in three average rarefied ZOTU tables to avoid the loss of samples with lower number of reads.

### 2.4. Statistical analysis

To analyze the influence of environmental gradients and the relative contribution of spatial versus temporal drivers on prokaryotic diversity on microbial communities, Cluster Analysis with Heatmap, Non-metric Multidimensional Scaling (nMDS) and Principal Coordinates Analysis (PCoA) were performed at the taxonomic level of ZOTUs based on Bray-Curtis dissimilarity (Legendre & Anderson, 1999). Heatmap workflows were carried out using the “pheatmap” package (version 1.0.12) in the R programming environment (Version 4.5.2 R Core Team, 2025) and PRIMER 7 to describe community dissimilarity in unconstrained space. ZOTU tables were square root transformed and standardized to totals (as recommended by Legendre & Gallagher, 2001). PCoA ordinations based on Bray-Curtis dissimilarities were performed using PRIMER 7 to describe the microbial community structure. For the environmental data PCoA, a matrix of environmental variables for the sampling points was square root transformed and normalized. The species matrix was composed of the ZOTU table for each sampling point and wetland. For the Heatmap analysis at the ZOTU level, the matrix comprised the most abundant ZOTUs (top 99% cumulative abundance). A univariate permutational analysis of variance (PERMANOVA) was performed with PRIMER 7; (Anderson, 2014) with 999 permutations was performed to analyze the effects of season, site, and subsite factors.

To identify the prokaryotic genera characteristic of each conservation status (well-preserved, altered, restored) within each Case Pilot, a Multilevel Pattern Analysis was conducted using the Indicator Value (IndVal) index proposed by Dufrêne and Legendre (1997). The analyses were performed using the multipatt function of the “*indicspecies*” package (ver. 1.7.9; De Cáceres et al., 2010) in the R statistical environment (Version 4.5.2 R Core Team, 2025). The “*IndVal*.g” variant of the index was specifically selected, which adjusts calculations to correct for the effect of unequal sizes among site groups (De Cáceres & Legendre, 2009). This index quantifies the indicator value based on the product of two independent components: specificity (component A), which represents the probability that a site belongs to the target group given the presence of the species; and fidelity (component B), which measures the frequency of occurrence of the species within the sites of that group. Following the methodology of De Cáceres et al. (2010), the analysis was not restricted to individual groups but evaluated all possible combinations of site categories (e.g., joint indicator species of altered and restored sites), allowing the detection of complex ecological patterns and shared niches. The statistical significance of the observed associations was determined using permutation tests (999 randomizations), selecting as valid indicators those genera with a p-value < 0.05 (Alonso et al., 2022).

## Results

### Seasonal patterns of the microbial community

Non-metric Multidimensional Scaling (nMDS) ordinations analysis, at ZOTU level, reveals that microbial community structure is primarily governed by environmental characteristics associated with a geographical/environmental component (Site) rather than seasonal variations in both water and sediment. Although a seasonal distribution is discernible within each site, the global ordination confirms that local environmental conditions act as the primary filter for community assembly over seasonal cycles. The tables showing relative abundance at the ZOTU level for water and sediment can be found in supplementary tables 1 and 2, respectively.

For the water samples, the nMDS ordination (Fig. 2A) reveals that biogeographical identity exerts a paramount influence on microbial community structure, clearly overriding seasonal signals. While intra-site temporal shifts are detectable, no cohesive global seasonal pattern emerges (Fig. 2B), indicating that local environmental filters are the primary drivers of assembly. The ordination space reveals a distinct gradient along the first axis: the marine-influenced systems of RI and the DU form a cohesive cluster, exhibiting high compositional similarity. At one extreme of the primary axis, the freshwater-dominated systems of the DA and CU converge, displaying closely related community profiles. Conversely, VA occupies the opposite extreme of this gradient, characterized by marked internal heterogeneity among its subsites, likely reflecting diverse hydrological management. CA occupies an intermediate position in the ordination, effectively bridging the divergent assemblages of VA and the continental DA-CU cluster.

**Figure 2.**
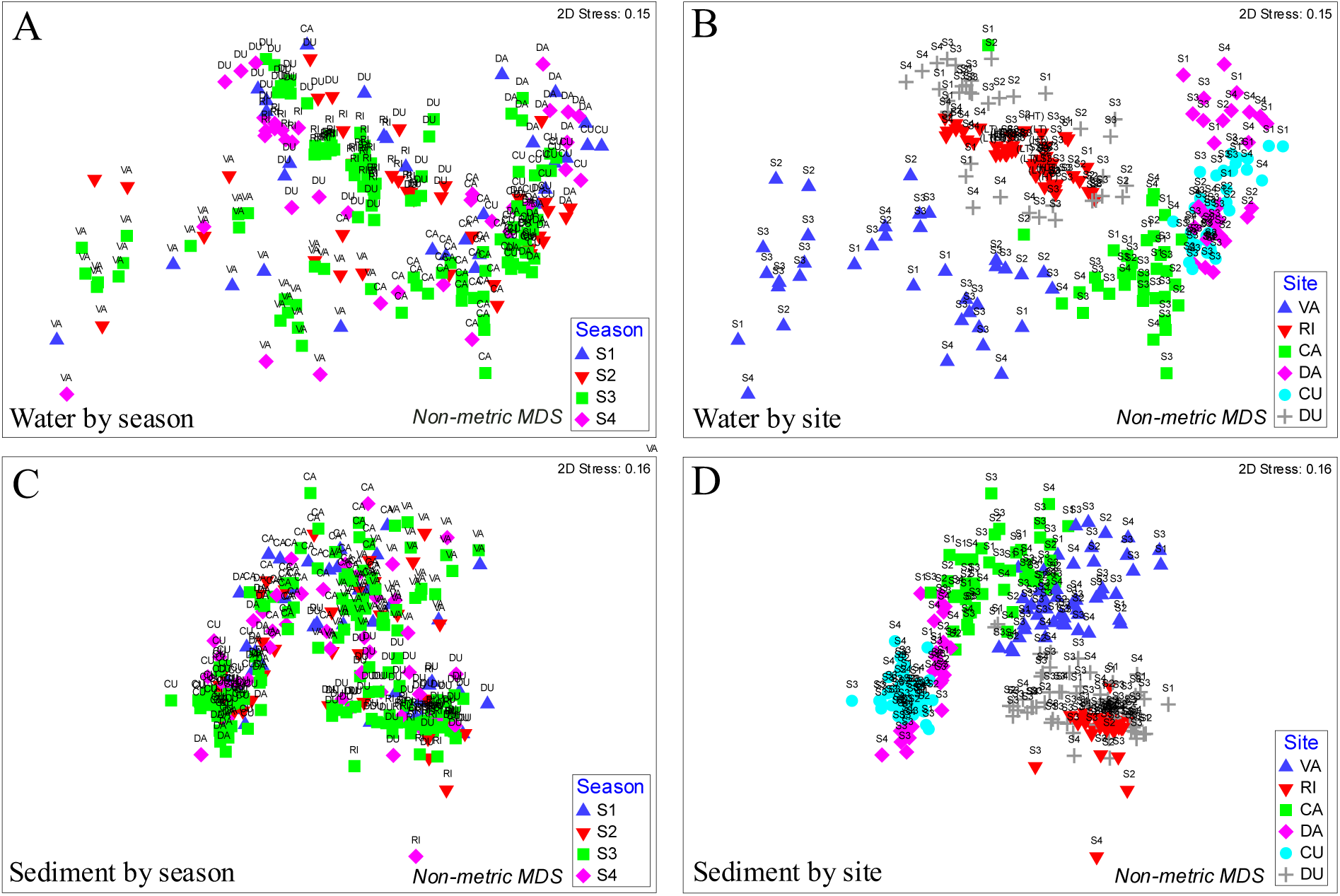
nMDS plots illustrate the separation of samples based upon differences in microbial community structure. A-B) Water samples grouped by site (A) and by season (B). C-D) Sediment samples grouped by site (C) and by season (D). Site (Camargue (CA), Curonian Lagoon (CU), Ria de Aveiro (RI), SW Dutch Delta (DU), Danube Delta (DA), Marjal dels Moros (VA)). Season (S1: Autumn, S2: Winter, S3: Spring, S4: Summer).

The sediment compartment mirrors this biogeographical structuring but with significantly reduced variability, confirming the higher inertia of microbioal communities in sediment compared to the water (Fig. 2C). Seasonality imposes no discernible global pattern, with samples grouping tightly by site regardless of the sampling season (Fig. 2D). The topological arrangement reinforces the gradient observed in the pelagic phase: the sediment communities of RI and DU are closely aligned, clearly distinguishing themselves from the cohesive continental block formed by DA and CU. VA again anchors the opposing end of the spectrum, though with tighter clustering than in water, while the CA serves as a transitional node connecting the Mediterranean and continental systems. This robust site-specific clustering confirms that edaphic and regional factors are the distinct determinants of sediment microbiome identity, buffering the community against the seasonal fluctuations that influence the overlying water.

PERMANOVA results (Table 2) revealed contrasting seasonal dynamics between the water and sediment matrices. In water, seasonality emerged as a determinant factor primarily in the CU and the DA. Specifically, in CU, significant differences were observed between most seasons within the altered and restored habitats (e.g., S1 vs S2, p=0.012 in A; S2 vs S3, p=0.001 in R), whereas in DA, seasonal variability was more pronounced in the well-preserved sites. Conversely, in ecosystems such as VA, CA, and DU, most seasonal comparisons in water were non-significant, indicating a in some cases temporally stable pelagic community or other drivers rather than temporal variability might be obscuring any potential seasonal trend. In the sediment matrix, temporal stability was even more generalized; most of the comparisons yielded non-significant values (p > 0.05) across sites like VA, CA, RI, and DU, regardless of habitat type. Exceptions were limited to CU, where well-preserved and restored habitats showed significant shifts, and sporadically in the well-preserved sites of DA, confirming that microbial community structure in sediment is considerably more resistant to seasonal fluctuations than that of the water.

**Table 2.** Pairwise PERMANOVA analysis with Monte Carlo correction assessing significant seasonal differences in prokaryotic community structure within each subsite. Case Pilot codes: CA (Camargue), CU (Curonian Lagoon), DA (Danube Delta), DU (Southwest Dutch Delta), RI (Ria de Aveiro), and VA (Marjal dels Moros). Subsite conservation status is denoted as: WP (Well-Preserved), A (Altered), and R (Restored). Seasonal campaigns are abbreviated as: S1 (Autumn), S2 (Winter), S3 (Spring), and S4 (Summer). Values represent statistical significance (p-value); p < 0.05 indicates significant differences between seasons. Asterisks indicate significance levels: *p < 0.05, **p < 0.01.

| *Water* | VA |  |  | CA |  |  | DA |  |  | CU |  |  | RI |  |  | DU |  |  |
| --- | --- | --- | --- | --- | --- | --- | --- | --- | --- | --- | --- | --- | --- | --- | --- | --- | --- | --- |
|  | A | WP | R | A | WP | R | A | WP | R | A | WP | R | A | WP | R | A | WP | R |
| *S1, S2* | 0.464 | 0.556 | 0.253 | 0.203 | 0.497 | 0.642 | n.d. | 0.061 | 0.565 | 0.012* | 0.125 | 0.085 | 0.367 | 0.252 | n.d. | 0.573 | 0.492 | 0.557 |
| *S1, S3* | 0.094 | 0.511 | 0.443 | 0.206 | 0.156 | 0.058 | 0.038* | 0.003* | 0.520 | 0.049* | 0.023* | 0.002* | 0.280 | 0.213 | n.d. | 0.242 | 0.256 | 0.322 |
| *S1, S4* | 0.490 | n.d. | 0.490 | 0.119 | 0.312 | 0.309 | n.d. | 0.130 | 0.544 | 0.193 | 0.514 | 0.299 | 0.173 | 0.212 | n.d. | 0.523 | 0.506 | 0.515 |
| *S2, S3* | 0.140 | 0.390 | 0.106 | 0.450 | 0.202 | 0.172 | 0.050* | 0.001* | 0.352 | 0.030* | 0.001* | 0.001* | 0.415 | 0.337 | 0.289 | 0.412 | 0.457 | 0.460 |
| *S2, S4* | 0.430 | 0.422 | 0.305 | 0.398 | 0.315 | 0.407 | n.d. | 0.047* | 0.284 | 0.189 | 0.042* | 0.144 | 0.092 | 0.094 | 0.029* | 0.485 | 0.513 | 0.527 |
| *S3, S4* | 0.037* | 0.274 | 0.343 | 0.158 | 0.098 | 0.174 | 0.015* | 0.004* | 0.228 | 0.093 | 0.007* | 0.005* | 0.131 | 0.074 | 0.033* | 0.419 | 0.300 | 0.298 |

| *Sediment* | VA |  |  | CA |  |  | DA |  |  | CU |  |  | RI |  |  | DU |  |  |
| --- | --- | --- | --- | --- | --- | --- | --- | --- | --- | --- | --- | --- | --- | --- | --- | --- | --- | --- |
|  | A | WP | R | A | WP | R | A | WP | R | A | WP | R | A | WP | R | A | WP | R |
| *S1, S2* | 0.632 | 0.570 | 0.622 | 0.506 | 0.419 | 0.544 | n.d. | n.d. | 0.597 | 0.690 | 0.432 | 0.229 | 0.499 | 0.554 | 0.646 | 0.661 | 0.569 | 0.381 |
| *S1, S3* | 0.645 | 0.645 | 0.517 | 0.566 | 0.710 | 0.470 | 0.420 | 0.015* | 0.482 | 0.211 | 0.042* | 0.004* | 0.495 | 0.426 | 0.651 | 0.394 | 0.428 | 0.403 |
| *S1, S4* | 0.563 | 0.607 | 0.614 | 0.486 | 0.561 | 0.575 | 0.382 | 0.307 | 0.444 | 0.454 | 0.518 | 0.190 | 0.592 | 0.510 | 0.540 | 0.590 | 0.605 | 0.646 |
| *S2, S3* | 0.684 | 0.489 | 0.692 | 0.402 | 0.366 | 0.709 | 0.623 | 0.013* | 0.633 | 0.147 | 0.048* | 0.003* | 0.656 | 0.470 | 0.693 | 0.462 | 0.307 | 0.500 |
| *S2, S4* | 0.634 | 0.608 | 0.720 | 0.635 | 0.509 | 0.428 | 0.568 | 0.276 | n.d. | 0.412 | 0.439 | 0.123 | 0.698 | 0.616 | 0.564 | 0.587 | 0.667 | 0.555 |
| *S3, S4* | 0.460 | 0.555 | 0.697 | 0.443 | 0.606 | 0.383 | 0.486 | 0.441 | 0.422 | 0.045* | 0.108 | 0.004* | 0.677 | 0.569 | 0.672 | 0.409 | 0.571 | 0.754 |

### Environmental factors and microbial community fingerprinting in spring

To identify the main biotic and abiotic drivers structuring the ecosystems studied during the period of maximum biological activity, a Principal Coordinates Analysis (PCoA) was performed on the physicochemical and biological variables of the water and sediment corresponding to the spring campaign (S3). This analysis enables the visualization of environmental variability across the six case pilots within a low-dimensional space, revealing how distinct hydrological, trophic, and management pressures shape the fundamental ecological niche during the time of highest annual productivity.

In the water (Figure 3), the two main axis of the ordination explains 44.3% of the total cumulative variation and reveals strong habitat structuring. The first axis (PCO1, 26%) establishes a clear dichotomy, drastically separating VA from the rest of the CPs. This segregation is driven by a unique hydrochemistry in VA, characterized by a saline gradient, a significant load of nitrogenous nutrients (NH_3_ and Total-N), as well as higher basal biomass, evidenced by high levels of bacterial abundance and chlorophyll-a (Chl-*a*). On the other hand, the second axis (PCO2, 18.3%) delineates a gradient separating the continental and eutrophic systems, such as DA and CU at the negative end, from the DU at the positive end, placing the CA and RI in an intermediate position, likely reflecting a transition in conditions of marine influence and phosphorus availability. In contrast, the analysis of sediment variables (Figure 3) shows less defined differentiation patterns among most sites, indicating greater homogeneity in the basal edaphic characteristics of the coastal wetlands studied. However, DA emerges as a notable exception, separating from the rest of the CPs along both main axes (PCO1 63.8% and PCO2 17.6%). This divergence suggests that the Danube sediment possesses distinctive physicochemical properties, possibly linked to its high rate of fluvial sedimentation, nutrient content, and organic matter, which functionally differentiate it from the more stable or saline substrates of the other coastal deltas and lagoons.

**Figure 3.**
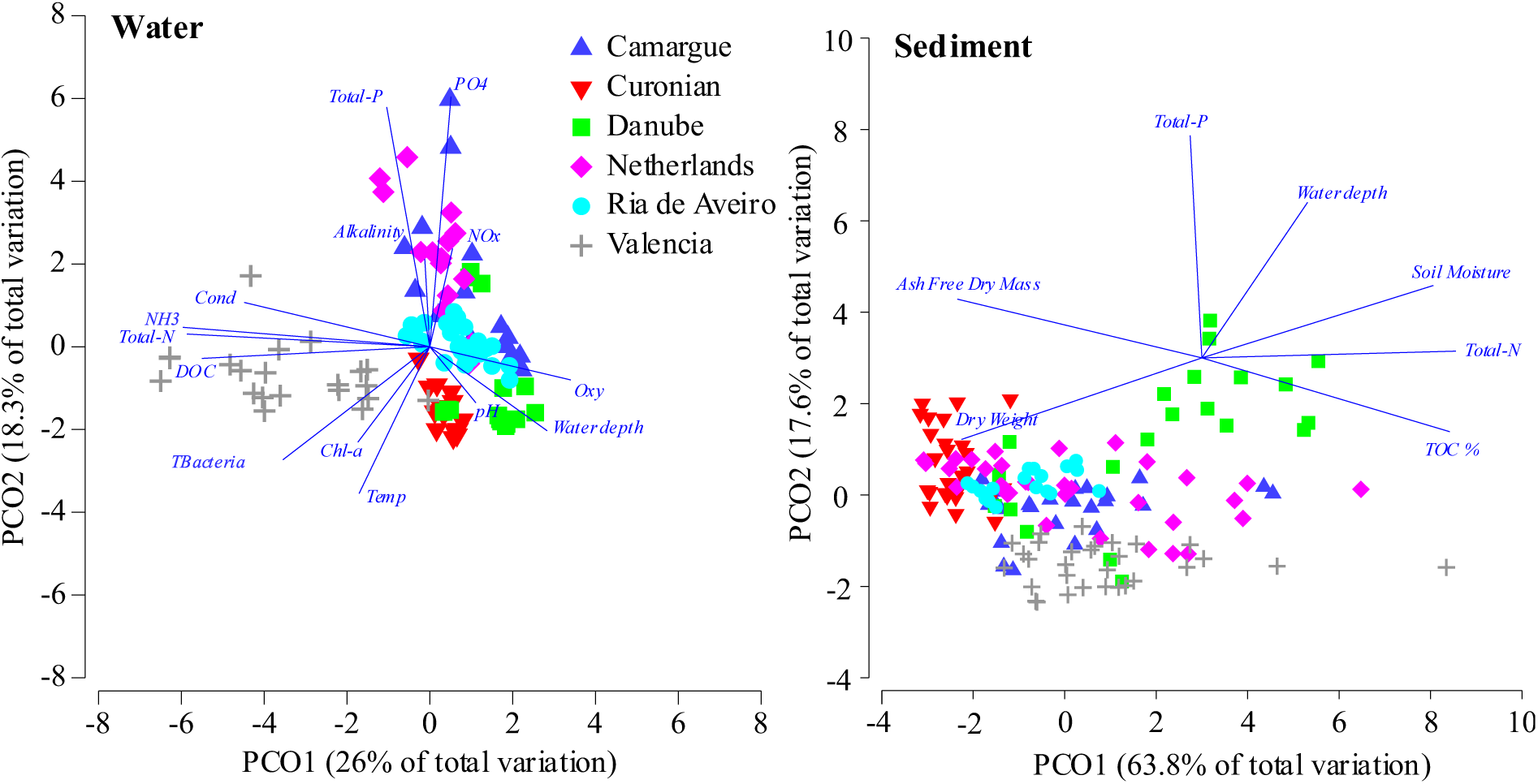
Principal Coordinates Analysis (PCoA) illustrates the environmental variability across the six European coastal wetland Case Pilots. The ordination is based on Euclidean distances calculated from normalized physicochemical variables measured in water and sediment. Vectors (arrows) indicate the direction and magnitude of the environmental parameters driving the separation between sites along the first two axes. Environmental Variables abbreviations: Temp: Temperature; Cond: Conductivity; Oxy: Dissolved oxygen; Chl-*a*: Chlorophyll-*a*; Tbacteria: Total bacterial abundance, DOC: Dissolved organic carbon, NH4: Ammonium, NOx: nitrate + nitrite, PO4: Ortophosphate, TN: Total Nitrogen; TP: Total Phosphorus; TOC: Total organic carbon.

Regarding the microbial community fingerprinting, heatmap analysis of ZOTUs representing 99% of the relative abundance in water samples (Figure 4), reveals the formation of blocks or clusters of distinctive ZOTUs for each geographic location, with very low taxonomic overlap between distant sites such as VA, CA, or the DA. This strong regional identity dominates the clustering hierarchy.

**Figure 4.**
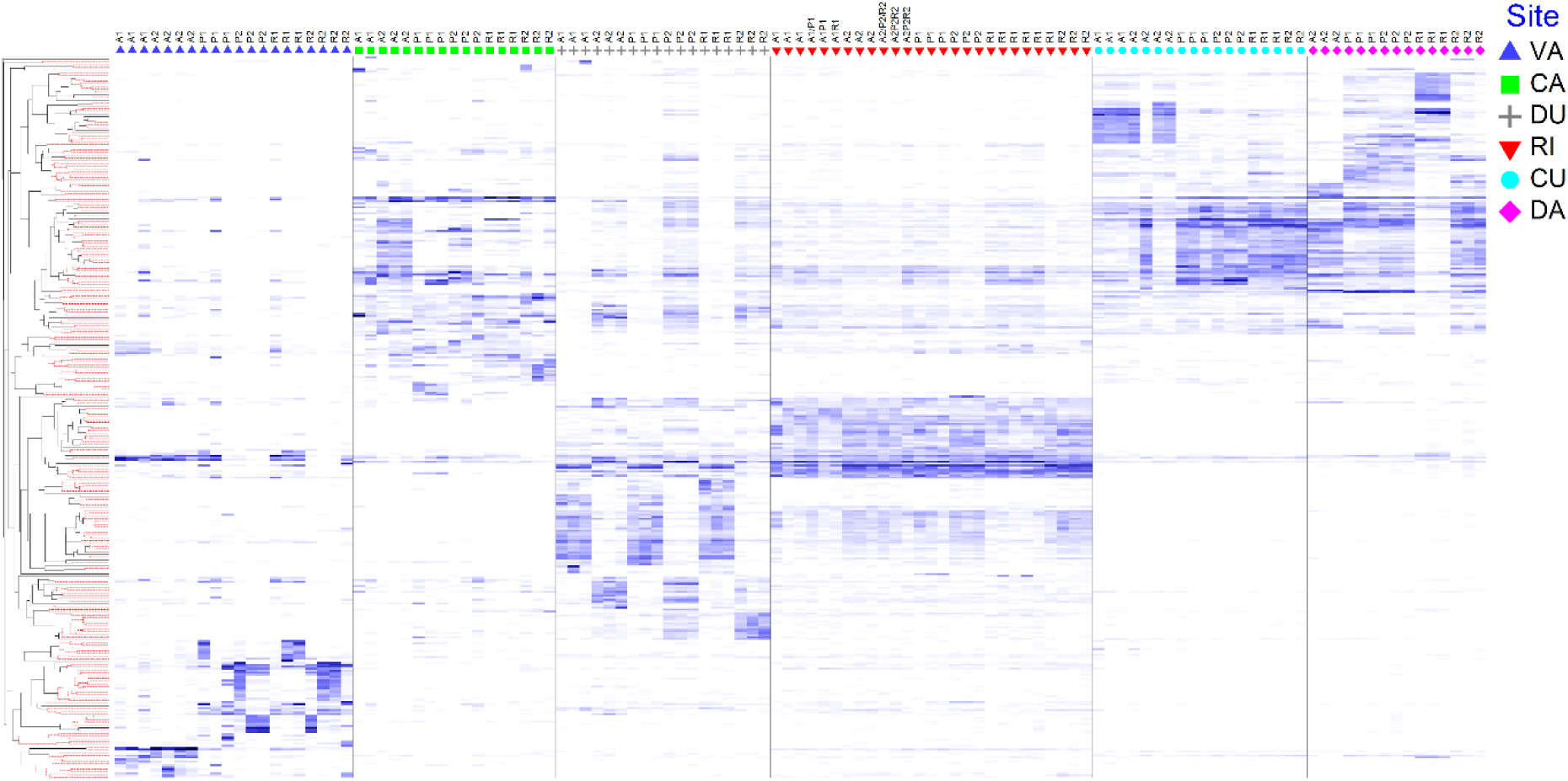
Heatmap analysis visualizing the relative abundance of the dominant prokaryotic ZOTUs across the study sites in water. The dataset was filtered to include only the ZOTUs contributing to the top 99% of total relative abundance to focus on the most prevalent taxa. Rows represent ZOTUs, clustered by Bray-Curtis co-occurrence patterns (left dendrogram). The color gradient indicates relative abundance (normalized Z-scores). CA: Camargue; CU: Curonian Lagoon; DA: Danube Delta; DU: Dutch Delta; RI: Ria de Aveiro; VA: Marjal dels Moros. Subsites: A1/A2: Altered; WP1/WP2: Well-Preserved; R1/R2: Restored.

This heatmap analysis (Figure 4) reveals a hierarchical community structure primarily defined by the water characteristics (Figure 3). At the regional scale, the DA and CU exhibit a remarkably similar microbial composition, forming a cohesive continental cluster. Similarly, the RI and the DU display significant compositional overlaps, likely reflecting shared estuarine characteristics, although each retains distinct, site-specific ZOTU clusters. CA occupies an intermediate position, showing partial affinities with the continental DA/CU block but maintaining a differentiated profile. Most notably, VA emerges as a clear outlier, characterized by a highly divergent community structure that bears little resemblance to the other ecosystems studied.

Embedded within these dominant geographic blocks, the fine-scale analysis reveals consistent compositional signatures driven by local management regimes (altered, well-preserved, restored). This is particularly visible in VA, the DU, and the CA, where distinct sub-clusters of ZOTUs emerge that are exclusively abundant in altered sites (A1/A2) but are effectively filtered out or severely depleted in well-preserved (WP1/WP2) and restored (R1/R2) habitats. Furthermore, a gradient of recovery is discernible across restoration stages; for instance, in the DU and RI, the ZOTU profiles of restored sites (R1/R2) do not form a monolithic block but instead display transitional assemblages that gravitate towards either well-preserved (WP) or altered (A) configurations depending on restoration maturity. This pattern demonstrates that while geography (environmental variables) establishes the regional species pool, habitat management exerts a critical secondary selective pressure that fine-tunes the local pelagic bacterial community assembly.

The sediment heatmap analysis (Figure 5), based on the ZOTUs contributing to 99% of the relative abundance, confirms an extremely robust site structuring, analogous to that observed in the water but with even more sharply defined dominance patterns. Dense and exclusive blocks of ZOTUs (high-abundance clusters) are identified as diagnostic for each study site, with minimal taxonomic overlap between geographic locations. This local specificity corroborates that local environmental and edaphic conditions are the determining drivers of sedimentc microbiome composition.

**Figure 5.**
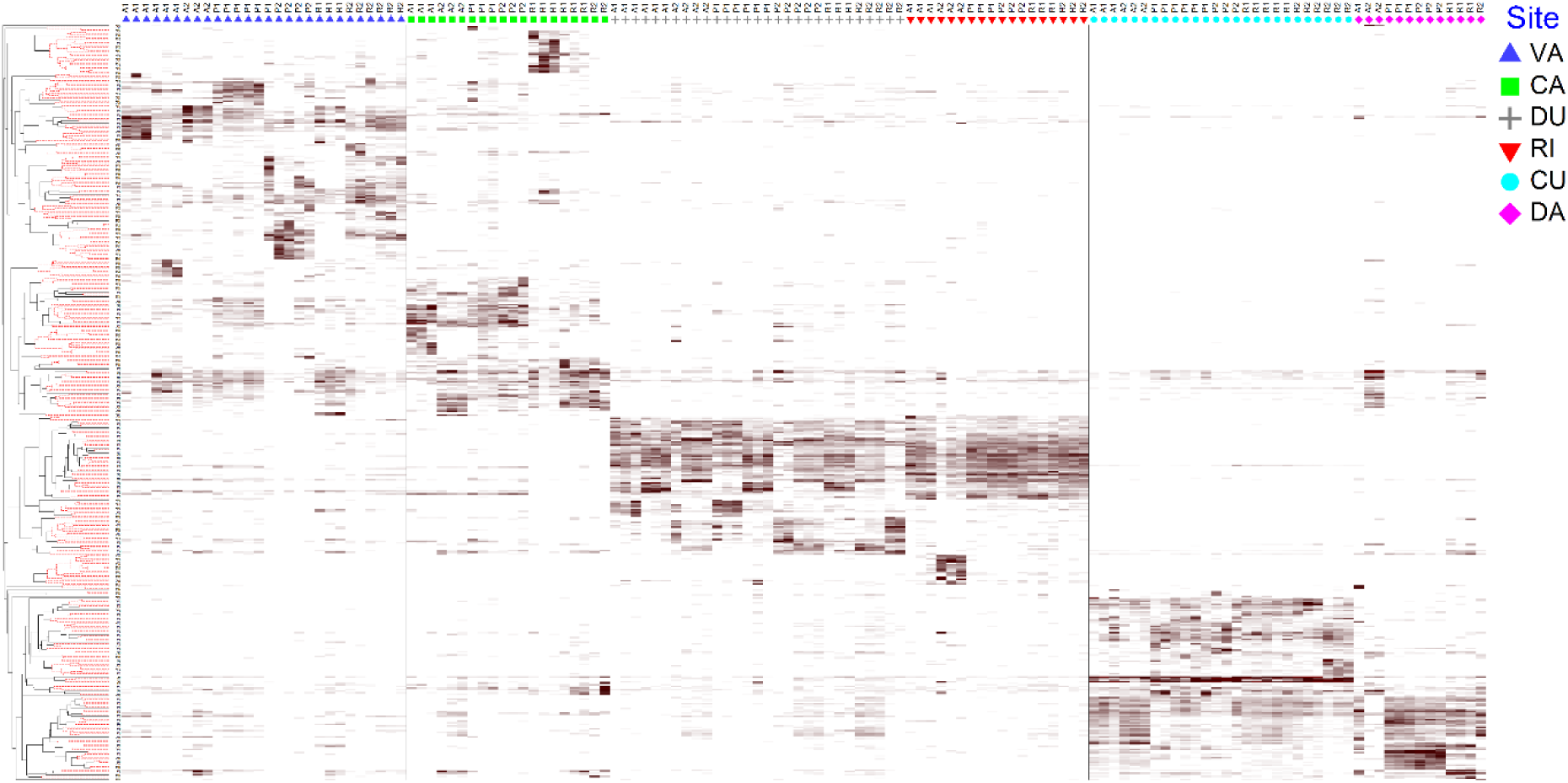
Heatmap analysis visualizing the relative abundance of the dominant prokaryotic ZOTUs across the study sites in the sediment. The dataset was filtered to include only the ZOTUs contributing to the top 99% of total relative abundance to focus on the most prevalent taxa. Rows represent ZOTUs, clustered by Bray-Curtis co-occurrence patterns (left dendrogram). The color gradient indicates relative abundance (normalized Z-scores). CA: Camargue; CU: Curonian Lagoon; DA: Danube Delta; DU: Dutch Delta; RI: Ria de Aveiro; VA: Marjal dels Moros. Subsites: A1/A2: Altered; WP1/WP2: Well-Preserved; R1/R2: Restored.

At the site level, DA and CU maintain clear similarities in microbial community composition, clustering within a broad continental group; however, CU distinguishes itself through specific species clades absent in DA, reflecting its unique biogeochemical identity. Similarly, RI and the DU display a comparable high-level composition, consistent with their Atlantic-influenced nature. In contrast, both CA and VA present highly specific ZOTU clades that are sharply differentiated from the rest of the sites. Heatmap shows intra-site variability, significant compositional variations dictated by conservation status. Within geographic blocks such as the DU and RI, specific ZOTU sub-clusters clearly differentiate altered sediments (A1/A2) from well-preserved ones (WP1/WP2). The dynamics of restored subsites are particularly notable; for instance, in DA and the DU, the profiles of restored subsites exhibit divergent behaviors, where some restored ZOTU clusters align with preserved patterns, indicating significant changes in community structure, while others retain taxonomic signatures shared with altered sites or present a unique transitional community.

The PERMANOVA analysis conducted for each site revealed heterogeneous patterns of differentiation among subsites in water (Table 3). In VA, significant structural differences were driven principally by the distinct composition of altered sites (A1 and A2) compared to the well-preserved (WP) and restored (R) ones, with no major significant variation observed between P and R, suggesting a successful convergence of the restored community towards the reference state. Conversely, in CA, differences among subsites were largely non-significant, indicating a high degree of homogeneity or connectivity across the conservation status gradient. DA presented a highly structured landscape where all subsites exhibited significantly different microbial compositions (p < 0.01), although P1, P2, and R2 shared a slightly higher degree of similarity (0.01 < p < 0.05). In CU, the main driver of dissimilarity was the altered site A1, which differed significantly from the rest of the locations. In RI, while the general pattern reflected a well-mixed estuarine system with low overall differentiation, the restored sites (R1 and R2) exhibited a notably similar microbial composition, clustering closely together. Finally, in DU, the analysis confirmed strong microbial community structure differences; specifically, the altered site A1 differed significantly from both the restored (R1) and well-preserved (WP1) sites.

**Table 3.** Statistical summary of the PERMANOVA pairwise comparisons analysis (Bray-Curtis distance, 999 permutations) for microbial community structure in water samples (spring samples S3) across different study sites. t: pairwise test statistic; P(MC): Monte Carlo statistical significance. Subsites groups: A1/A2: Altered; WP1/WP2: Well-Preserved; R1/R2: Restored. Asterisks indicate significance levels: *p < 0.05, **p < 0.01.

| Within level 'VA' of factor 'Site' |  |  |  |  | Within level 'CA' of factor 'Site' |  |  |  |  | Within level 'DA' of factor 'Site' |  |  |  |  |
| --- | --- | --- | --- | --- | --- | --- | --- | --- | --- | --- | --- | --- | --- | --- |
| Groups | t | P(perm) | Unique perms | P(MC) | Groups | t | P(perm) | Unique perms | P(MC) | Groups | t | P(perm) | Unique perms | P(MC) |
| A1, A2 | 1.742 | 0.062 | 35 | 0.053 | A1, A2 | 1.566 | 0.094 | 10 | 0.134 | A1, A2 | No test |  |  |  |
| A1, R1 | 1.926 | 0.099 | 10 | 0.036* | A1, R1 | 1.811 | 0.087 | 10 | 0.081 | A1, R1 | No test |  |  |  |
| A1, R2 | 2.051 | 0.022 | 35 | 0.017* | A1, R2 | 1.280 | 0.104 | 10 | 0.245 | A1, R2 | No test |  |  |  |
| A1, WP1 | 1.568 | 0.083 | 10 | 0.069 | A1, WP1 | 1.443 | 0.106 | 10 | 0.149 | A1, WP1 | No test |  |  |  |
| A1, WP2 | 2.316 | 0.118 | 10 | 0.016* | A1, WP2 | 1.025 | 0.492 | 10 | 0.427 | A1, WP2 | No test |  |  |  |
| A2, R1 | 2.315 | 0.029 | 35 | 0.015* | A2, R1 | 2.017 | 0.092 | 10 | 0.044* | A2, R1 | 6.645 | 0.107 | 10 | 0.001** |
| A2, R2 | 2.586 | 0.021 | 35 | 0.003** | A2, R2 | 1.675 | 0.097 | 10 | 0.076 | A2, R2 | 3.040 | 0.106 | 10 | 0.006** |
| A2, WP1 | 1.925 | 0.025 | 35 | 0.024* | A2, WP1 | 1.879 | 0.095 | 10 | 0.046* | A2, WP1 | 4.007 | 0.108 | 10 | 0.003** |
| A2, WP2 | 2.971 | 0.032 | 35 | 0.006** | A2, WP2 | 1.558 | 0.109 | 10 | 0.105 | A2, WP2 | 4.099 | 0.102 | 10 | 0.001** |
| R1, R2 | 1.615 | 0.024 | 35 | 0.045* | R1, R2 | 1.750 | 0.098 | 10 | 0.066 | R1, R2 | 4.971 | 0.090 | 10 | 0.003** |
| R1, WP1 | 0.710 | 0.803 | 10 | 0.671 | R1, WP1 | 2.188 | 0.114 | 10 | 0.03* | R1, WP1 | 5.225 | 0.110 | 10 | 0.002** |
| R1, WP2 | 1.841 | 0.103 | 10 | 0.057 | R1, WP2 | 1.697 | 0.095 | 10 | 0.067 | R1, WP2 | 6.241 | 0.099 | 10 | 0.001** |
| R2, WP1 | 1.261 | 0.148 | 35 | 0.211 | R2, WP1 | 1.753 | 0.107 | 10 | 0.053 | R2, WP1 | 2.646 | 0.117 | 10 | 0.010* |
| R2, WP2 | 0.853 | 0.375 | 35 | 0.514 | R2, WP2 | 1.337 | 0.102 | 10 | 0.181 | R2, WP2 | 2.922 | 0.099 | 10 | 0.011* |
| WP1, WP2 | 1.471 | 0.108 | 10 | 0.118 | WP1, WP2 | 1.497 | 0.104 | 10 | 0.106 | WP1, WP2 | 2.257 | 0.093 | 10 | 0.024* |

| Within level 'CU' of factor 'Site' |  |  |  |  | Within level 'RI' of factor 'Site' |  |  |  |  | Within level 'DU' of factor 'Site' |  |  |  |  |
| --- | --- | --- | --- | --- | --- | --- | --- | --- | --- | --- | --- | --- | --- | --- |
| Groups | t | P(perm) | Unique perms | P(MC) | Groups | t | P(perm) | Unique perms | P(MC) | Groups | t | P(perm) | Unique perms | P(MC) |
| A1, A2 | 1.105 | 0.320 | 35 | 0.314 | A1, A2 | 1.377 | 0.217 | 10 | 0.163 | A1, A2 | 2.592 | 0.103 | 10 | 0.014* |
| A1, R1 | 5.384 | 0.102 | 10 | 0.001** | A1, R1 | 1.190 | 0.180 | 84 | 0.247 | A1, R1 | 1.547 | 0.107 | 10 | 0.119 |
| A1, R2 | 4.679 | 0.105 | 10 | 0.006** | A1, R2 | 1.763 | 0.116 | 10 | 0.069 | A1, R2 | 3.192 | 0.093 | 10 | 0.004** |
| A1, WP1 | 3.309 | 0.101 | 10 | 0.004** | A1, WP1 | 1.204 | 0.322 | 10 | 0.273 | A1, WP1 | 1.106 | 0.190 | 10 | 0.319 |
| A1, WP2 | 4.750 | 0.104 | 10 | 0.002** | A1, WP2 | 1.870 | 0.100 | 10 | 0.047* | A1, WP2 | 3.823 | 0.093 | 10 | 0.005** |
| A2, R1 | 2.113 | 0.063 | 35 | 0.054 | A2, R1 | 2.108 | 0.027 | 84 | 0.019* | A2, R1 | 2.645 | 0.087 | 10 | 0.014* |
| A2, R2 | 1.703 | 0.279 | 15 | 0.127 | A2, R2 | 1.527 | 0.116 | 10 | 0.117 | A2, R2 | 2.274 | 0.098 | 10 | 0.012* |
| A2, WP1 | 1.805 | 0.114 | 35 | 0.069 | A2, WP1 | 1.620 | 0.095 | 10 | 0.095 | A2, WP1 | 2.577 | 0.096 | 10 | 0.011* |
| A2, WP2 | 2.156 | 0.052 | 35 | 0.043* | A2, WP2 | 1.640 | 0.086 | 10 | 0.092 | A2, WP2 | 1.939 | 0.084 | 10 | 0.036* |
| R1, R2 | 2.078 | 0.099 | 10 | 0.062 | R1, R2 | 2.398 | 0.010 | 84 | 0.004** | R1, R2 | 3.282 | 0.105 | 10 | 0.009 |
| R1, WP1 | 2.037 | 0.091 | 10 | 0.046* | R1, WP1 | 1.541 | 0.054 | 84 | 0.078 | R1, WP1 | 1.565 | 0.095 | 10 | 0.082 |
| R1, WP2 | 3.452 | 0.117 | 10 | 0.005** | R1, WP2 | 2.451 | 0.019 | 84 | 0.01* | R1, WP2 | 3.983 | 0.100 | 10 | 0.005 |
| R2, WP1 | 1.369 | 0.199 | 10 | 0.220 | R2, WP1 | 1.811 | 0.096 | 10 | 0.065 | R2, WP1 | 3.156 | 0.101 | 10 | 0.007 |
| R2, WP2 | 2.646 | 0.102 | 10 | 0.031* | R2, WP2 | 1.790 | 0.093 | 10 | 0.061 | R2, WP2 | 2.635 | 0.104 | 10 | 0.009 |
| WP1, WP2 | 1.068 | 0.595 | 10 | 0.334 | WP1, WP2 | 1.872 | 0.103 | 10 | 0.049 | WP1, WP2 | 3.811 | 0.098 | 10 | 0.004 |

In the sediment compartment (Table 4), PERMANOVA results highlighted site-specific patterns of microbial sediment community differentiation. In VA, a generalized structural divergence was observed, with most subsites presenting significantly different microbial compositions; notably, the highest similarities were found between A2 and R1, as well as between R2 and P2, suggesting specific localized recovery trajectories. In CA, sediment communities exhibited higher homogeneity, with no significant differences detected across several comparisons (A1 vs. R2, WP1, WP2; A2 vs. R2; R1 vs. R2; WP1 vs. WP2; and R2 vs. WP1, WP2), while the remaining pairwise contrasts showed moderate differentiation (0.01 < p < 0.05). DA displayed a unique pattern where the altered site A1 did not differ significantly from any other subsite, nor did the pairs WP1-WP2, A2-R2, and R1-R2; however, other comparisons revealed significant shifts (0.01 < p < 0.05). Similarly, in CU, A1 showed no significant differences with the rest of the subsites, mirroring the lack of differentiation between R2 and the well-preserved sites (WP1, WP2) as well as between P1 and P2 themselves; the primary drivers of dissimilarity here were the contrasts between A2 and the restored sites (R1, R2). In RI, the altered site A2 emerged as the distinct outlier, showing significant differences (0.01 < p < 0.05) compared to the rest of the subsites, which otherwise shared a homogeneous structure. Finally, in DU, the strongest significant differences (p < 0.01) were driven by the divergence of A1 from the mature restored (R2) and well-preserved (WP2) sites; to a lesser extent (0.01 < p < 0.05), differentiation was also observed between A2 and R1, as well as in comparisons involving R1 (vs. R2, P2) and P1 (vs. R2, P2), reflecting a gradient of recovery maturity.

**Table 4.** Statistical summary of the PERMANOVA pairwise comparisons analysis (Bray-Curtis distance, 999 permutations) for microbial community structure in the sediment (spring samples S3) across the different study sites. t: pairwise test statistic; P(MC): Monte Carlo statistical significance. Subsites groups: A1/A2: Altered; WP1/WP2: Well-Preserved; R1/R2: Restored. Asterisks indicate significance levels: *p < 0.05, **p < 0.01.

| Within level 'VA' of factor 'Site' |  |  |  |  | Within level 'CA' of factor 'Site' |  |  |  |  | Within level 'DA' of factor 'Site' |  |  |  |  |
| --- | --- | --- | --- | --- | --- | --- | --- | --- | --- | --- | --- | --- | --- | --- |
| Groups | t | P(perm) | Unique perms | P(MC) | Groups | t | P(perm) | Unique perms | P(MC) | Groups | t | P(perm) | Unique perms | P(MC) |
| A1, A2 | 1.623 | 0.035 | 84 | 0.046 | A1, A2 | 2.313 | 0.105 | 10 | 0.019* | A1, A2 | 2.418 | 0.335 | 3 | 0.186 |
| A1, R1 | 1.623 | 0.023 | 84 | 0.049* | A1, R1 | 1.759 | 0.011 | 84 | 0.019* | A1, R1 | 1.502 | 0.233 | 4 | 0.198 |
| A1, R2 | 1.694 | 0.004 | 419 | 0.010* | A1, R2 | 1.627 | 0.105 | 10 | 0.101 | A1, R2 | No test |  |  |  |
| A1, WP1 | 1.630 | 0.006 | 418 | 0.018* | A1, WP1 | 1.391 | 0.081 | 10 | 0.148 | A1, WP1 | 2.528 | 0.250 | 4 | 0.076 |
| A1, WP2 | 1.794 | 0.008 | 209 | 0.021* | A1, WP2 | 1.606 | 0.108 | 10 | 0.080 | A1, WP2 | 2.308 | 0.229 | 4 | 0.079 |
| A2, R1 | 1.707 | 0.115 | 10 | 0.066 | A2, R1 | 1.738 | 0.030 | 84 | 0.019* | A2, R1 | 2.498 | 0.096 | 10 | 0.026* |
| A2, R2 | 1.993 | 0.012 | 84 | 0.008** | A2, R2 | 1.742 | 0.083 | 10 | 0.089 | A2, R2 | 2.229 | 0.305 | 3 | 0.172 |
| A2, WP1 | 1.866 | 0.013 | 84 | 0.012* | A2, WP1 | 1.943 | 0.104 | 10 | 0.036* | A2, WP1 | 3.655 | 0.096 | 10 | 0.012* |
| A2, WP2 | 2.028 | 0.024 | 35 | 0.017* | A2, WP2 | 2.039 | 0.099 | 10 | 0.032* | A2, WP2 | 3.434 | 0.104 | 10 | 0.014* |
| R1, R2 | 1.900 | 0.019 | 84 | 0.006** | R1, R2 | 1.185 | 0.100 | 28 | 0.254 | R1, R2 | 1.636 | 0.253 | 4 | 0.166 |
| R1, WP1 | 1.794 | 0.020 | 84 | 0.019* | R1, WP1 | 1.565 | 0.018 | 84 | 0.044* | R1, WP1 | 2.395 | 0.104 | 10 | 0.015* |
| R1, WP2 | 1.931 | 0.027 | 35 | 0.026* | R1, WP2 | 1.668 | 0.018 | 84 | 0.036* | R1, WP2 | 2.387 | 0.089 | 10 | 0.017* |
| R2, WP1 | 1.824 | 0.011 | 408 | 0.014* | R2, WP1 | 1.452 | 0.090 | 10 | 0.150 | R2, WP1 | 2.724 | 0.234 | 4 | 0.049* |
| R2, WP2 | 1.502 | 0.009 | 210 | 0.051 | R2, WP2 | 1.601 | 0.112 | 10 | 0.120 | R2, WP2 | 2.659 | 0.250 | 4 | 0.043* |
| WP1, WP2 | 1.840 | 0.009 | 208 | 0.016* | WP1, WP2 | 1.440 | 0.101 | 10 | 0.122 | WP1, WP2 | 1.843 | 0.084 | 10 | 0.061 |

| Within level 'CU' of factor 'Site' |  |  |  |  | Within level 'RI' of factor 'Site' |  |  |  |  | Within level 'DU' of factor 'Site' |  |  |  |  |
| --- | --- | --- | --- | --- | --- | --- | --- | --- | --- | --- | --- | --- | --- | --- |
| Groups | t | P(perm) | Unique perms | P(MC) | Groups | t | P(perm) | Unique perms | P(MC) | Groups | t | P(perm) | Unique perms | P(MC) |
| A1, A2 | 1.730 | 0.101 | 10 | 0.064 | A1, A2 | 1.748 | 0.108 | 10 | 0.046* | A1, A2 | 1.750 | 0.007 | 209 | 0.022* |
| A1, R1 | 1.583 | 0.012 | 56 | 0.063 | A1, R1 | 1.296 | 0.110 | 10 | 0.191 | A1, R1 | 1.371 | 0.028 | 84 | 0.099 |
| A1, R2 | 1.554 | 0.024 | 84 | 0.065 | A1, R2 | 1.455 | 0.103 | 10 | 0.138 | A1, R2 | 2.071 | 0.004 | 407 | 0.007** |
| A1, WP1 | 1.429 | 0.042 | 84 | 0.098 | A1, WP1 | 1.185 | 0.095 | 10 | 0.267 | A1, WP1 | 1.013 | 0.332 | 406 | 0.392 |
| A1, WP2 | 1.695 | 0.108 | 10 | 0.077 | A1, WP2 | 1.246 | 0.184 | 10 | 0.254 | A1, WP2 | 2.006 | 0.006 | 413 | 0.006** |
| A2, R1 | 2.225 | 0.012 | 56 | 0.008** | A2, R1 | 1.977 | 0.092 | 10 | 0.030* | A2, R1 | 1.736 | 0.028 | 35 | 0.046* |
| A2, R2 | 2.364 | 0.010 | 84 | 0.002** | A2, R2 | 2.055 | 0.100 | 10 | 0.028* | A2, R2 | 1.327 | 0.168 | 126 | 0.161 |
| A2, WP1 | 1.990 | 0.009 | 84 | 0.010* | A2, WP1 | 2.062 | 0.102 | 10 | 0.032* | A2, WP1 | 1.499 | 0.028 | 209 | 0.055 |
| A2, WP2 | 2.314 | 0.094 | 10 | 0.024* | A2, WP2 | 1.968 | 0.104 | 10 | 0.039* | A2, WP2 | 1.144 | 0.189 | 126 | 0.264 |
| R1, R2 | 1.747 | 0.003 | 423 | 0.032* | R1, R2 | 1.750 | 0.093 | 10 | 0.060 | R1, R2 | 1.900 | 0.018 | 56 | 0.026* |
| R1, WP1 | 1.729 | 0.005 | 395 | 0.035* | R1, WP1 | 1.281 | 0.105 | 10 | 0.187 | R1, WP1 | 1.330 | 0.048 | 84 | 0.140 |
| R1, WP2 | 1.843 | 0.017 | 56 | 0.021* | R1, WP2 | 1.481 | 0.107 | 10 | 0.114 | R1, WP2 | 1.868 | 0.019 | 56 | 0.031* |
| R2, WP1 | 1.479 | 0.036 | 411 | 0.066 | R2, WP1 | 1.162 | 0.409 | 10 | 0.296 | R2, WP1 | 1.829 | 0.004 | 411 | 0.011* |
| R2, WP2 | 1.496 | 0.046 | 84 | 0.079 | R2, WP2 | 0.901 | 0.821 | 10 | 0.523 | R2, WP2 | 0.964 | 0.512 | 125 | 0.404 |
| WP1, WP2 | 1.341 | 0.122 | 84 | 0.145 | WP1, WP2 | 0.881 | 0.694 | 10 | 0.544 | WP1, WP2 | 1.756 | 0.004 | 408 | 0.014* |

The Principal Coordinates Analysis (PCoA) based on ZOTU abundance in water samples (Figure 6) explained 48.7% of the total cumulative variation (32.5% on the PCO1 axis and 16.2% on the PCO2 axis). The ordination reveals a strong biogeographical structuring, where the first axis (PCO1) clearly separates systems with greater marine influence, RI and DU, located in the positive zone, from systems with more continental or brackish characteristics, such as DA, CU, and CA, in the negative zone. The second axis (PCO2) distinctively segregates VA from the rest of the ecosystems, isolating it at the bottom of the plot. The overlay of similarity contours (Bray-Curtis) reinforces that samples cluster primarily by geographic location (with intra-site similarities exceeding 20-40%), indicating that regional identity prevails over local management factors (altered, well-preserved, restored) in shaping the pelagic bacterial community structure.

**Figure 6.**
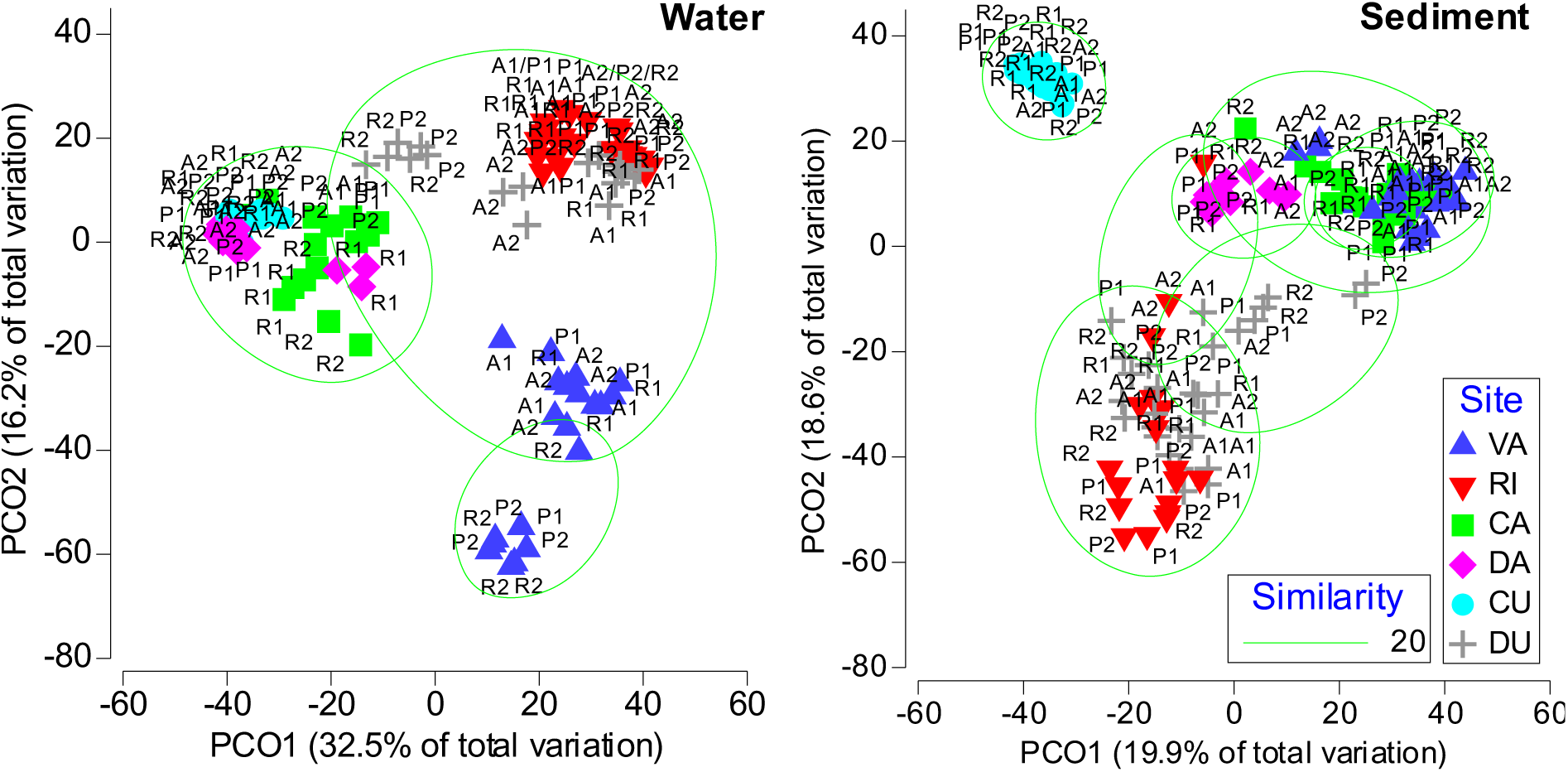
Principal Coordinates Analysis (PCoA) illustrating the microbial community variability across the six European coastal wetland Case Pilots. The ordination is based on Bray-Curtis distances calculated from standardized microbial composition (complete ZOTU table) measured in water and sediment. Subsites: A1/A2: Altered; WP1/WP2: Well-Preserved; R1/R2: Restored.

The Principal Coordinates Analysis (PCoA) for the sediment microbial community (Figure 6) explained 38.5% of the total variation (19.9% on PCO1 and 18.6% on PCO2). The ordination highlights the biogeochemical singularity of CU, which diverges drastically from the rest of the sites along the combination of PCO1 and PCO2 axis, forming an isolated cluster in the upper part of the graph. On the PCO1 axis, a continuous and well-defined biogeographical gradient is observed ordering the remaining ecosystems: the gradient begins at the positive end with VA, transitions through the intermediate systems of CA and DA, passes through DU, and culminates at the negative end with RI. This linear arrangement suggests a gradual transition in microbial community composition in sediment, possibly linked to a latitudinal or salinity gradient connecting Mediterranean to Atlantic systems, with VA and RI representing the opposite ends of this spectrum.

Comparing the ordinations between compartments reveals a fundamental shift in ecological differentiation patterns. While for water, VA is the most divergent ecosystem, clearly segregating from the rest along the PCO2 axis, likely due to specific local hydrological conditions. In the sediment, VA loses that exclusivity and integrates as one end of a biogeographical continuum. Conversely, CU, which is grouped closely with the DA in the water samples, emerges in the sediment as the most distinctive and isolated system. This indicates a decoupling of the drivers structuring both communities: water responds to a strong dichotomy where VA is unique, whereas the sediment reflects a smoother gradient between marine and Mediterranean sites, disrupted only by the singular sediment conditions of CU.

### Indicator Species Analysis

The IndVal analysis revealed marked differences in bacterial community structure among conservation status, identifying specific bioindicators for altered, restored, and well-preserved conditions. The tables showing relative abundance at the genus level can be found in supplementary tables 3 (water) and 4 (sediment), while the complete tables from the IndVal analysis can be found in supplementary tables 5 (water) and 6 (sediment).

#### Curonian Lagoon

In Curonian Lagoon, the IndVal analysis revealed a strong community structuring (p < 0.05) that clearly differentiates the dynamics between water (77 indicator species) and the sediment (117 species). In the water, altered zones (A1+A2) are defined by signals of anoxia and eutrophication, highlighting indicators of methane and sulfide production such as *Methyloparacoccus* (stat=0.98, p=0.005) and *Desulfatiglans* (stat=0.94, p=0.015), as well as a latent risk of algal blooms evidenced by the cyanobacterium *Dolichospermum* (stat=1.0, p=0.003). Conversely, well-preserved sites (WP1+WP2) exhibit the oligotrophic “target state,” dominated by ultramicrobacteria such as *Ca. Planktoluna* (stat=0.85, p=0.002) and *Rhodoluna* (stat=0.83, p=0.008). Restored sites (R1+R2) are in a successful but incomplete transition phase; they have effectively eliminated degradation indicators, yet instead of recruiting the preserved microbiota, they are dominated by freshwater generalists like *Sediminibacterium* (stat=0.84, p=0.001) and *Fluviicola* (stat=0.75, p=0.023). Unlike water, the sediment shows a functional divergence marked by “ecological memory.” Altered zones (A1+A2) function as intense anaerobic reactors, characterized by methanotrophs such as *Methyloglobulus* (stat=0.97, p=0.001) and *Methylococcus* (stat=0.90, p=0.003). Restored sites (R1) retain a strong imprint of this alteration, sharing a cluster with altered zones where *Methyloparacoccus* (stat=0.97, p=0.001) and *Dolichospermum* (stat=0.88, p=0.004) persist, suggesting that the sediment acts as a reservoir maintaining eutrophication potential. Nevertheless, restoration has fostered the development of new exclusive functions based on iron reduction, indicated by *Geomonas* (stat=0.96, p=0.001), differentiating them from the sulfur-driven processes of altered sites, whereas the well-preserved (WP) is distinguished by greater efficiency in the nitrogen cycle (*Ca. Nitrotoga*) and the presence of deep-biosphere archaea (*Hadarchaeum*).

#### Camargue

In Camargue, the water analysis revealed a strong community structuring (99 significant indicators, p<0.05) reflecting the management history. The altered sites (A1+A2) evidence the agricultural legacy through strict anaerobic bacteria such as *Thermoanaerobaculum* (stat=0.92, p=0.018) and *Syntrophorhabdus* (stat=0.89, p=0.018), typical of flooded and compacted soils, along with *Mesorhizobium* (stat=0.98, p=0.003) associated with crops. Following reconstruction, the restored sites (R1+R2) show a functional recovery marked by the reactivation of the sulfur cycle, indicated by *Thiovirga* (stat=0.97, p=0.007) and *Chromatium* (stat=1.0, p=0.009), and the presence of complex degraders like *Polyangium* (stat=0.94, p=0.010). Conversely, the well-preserved sites (WP1+WP2) host a mature and diverse ecosystem, characterized by greater saline influence and methylotrophic activity, with key indicators like *Halomicroarcula* (stat=1.0, p=0.006) and *Methylophaga* (stat=1.0, p=0.002). The analysis of sediment reveals a distinct dynamic, characterized by high ecological inertia and overlap between conservation status. The altered sites (A) retain a strong anoxic footprint dominated by fermenters like *Propionivibrio* (stat=0.94, p=0.001) and methanotrophs like *Methylocystis* (stat=0.96, p=0.003), signaling a persistent reducing environment. However, the restored sites (R) exhibit a functional soil recovery, diversifying towards the nitrogen cycle with root symbionts such as *Bradyrhizobium* (stat=0.95, p=0.001) and contaminant degradation with *Sphingopyxis* (stat=1.0, p=0.006). Finally, the well-preserved sites (WP) present a stable microbiome typical of mature soils, defined by specific indicators like *Blastocatella* (stat=0.95, p=0.002) and *Filomicrobium* (stat=1.0, p=0.004).

#### Marjal dels Moros

In Marjal dels Moros, water analysis (97 indicators, p<0.05) revealed a sharp dichotomy between the freshwater dominated altered sites and the restored sites with saline conditions. The altered sites (A) reflect the footprint of “freshening” and eutrophication through exclusive indicators of anoxia and the sulfur cycle such as *Solidesulfovibrio* (stat=1.0, p=0.004) and *Desulfomonile* (stat=0.95, p=0.001), along with cellulose fermenters like *Clostridium* (stat=0.89, p=0.002), confirming severe trophic alteration. Conversely, the restored sites (R) show a notable convergence with the well-preserved sites (WP), evidencing the success of hydrological restoration by recovering halophilic taxa such as *Salinivibrio* (p=0.001) and *Marinobacter* (p=0.025), which displace the freshwater microbiota. Nevertheless, R sites maintain a distinct signal of incipient remediation indicated by genera such as *Shinella* (stat=0.87, p=0.015). The sediment analysis displays greater complexity (289 indicators) and a slower recovery. The altered sites (A) retain an anaerobic and sulfurous legacy dominated by *Treponema* (stat=0.91, p=0.003) and sulfur cycle bacteria such as *Desulfococcus* (p=0.011) and *Thiocapsa* (p=0.001). Restoration has triggered a diversification in the restored sites (R), which, although retaining some inherited organic load, mark a transition towards marine conditions with indicators such as *Sulfitobacter* (stat=1.0, p=0.001), *Vibrio* (p=0.001), and biofilm formers like *Caulobacter* (p=0.001). Finally, the well-preserved Sites (WP) define the hypersaline reference target with strict specialists like *Imperialibacter* (stat=0.99, p=0.001) and *Desulfovermiculus* (stat=0.85, p=0.003).

#### Danube Delta

In the Danube Delta, water analysis (105 indicators, p<0.05) validates the success of hydrological reconnection through a clear ecological convergence between restored and well-preserved sites. The altered sites (A2) present a perturbed community with 17 exclusive indicators, where genera such as *Shewanella* (stat=1.0, p=0.008) and *Fusibacter* (stat=1.0, p=0.008) signal rapid anoxic conditions due to recent flooding, while *Arcobacter* (p=0.026) and *Cloacibacterium* (p=0.023) reveal fecal/organic contamination derived from prior agricultural and livestock use. In contrast, the well-preserved sites (WP) maintain a clean-water core characterized by *Polynucleobacter*, *Limnohabitans*, and cyanobacteria such as *Microcystis* (p=0.003). The restored sites (R) show successful “renaturalization,” sharing robust indicators with P such as *Roseomonas* (p=0.008) and a guild of picocyanobacteria including *Cyanobium* (stat=1.0, p=0.005), *Synechocystis* (stat=1.0, p=0.005), and *Geminocystis* (stat=1.0, p=0.007 in R1), that indicate stable, low-turbidity conditions and a phytoplankton community not dominated by bloom-forming cyanobacteria. Nevertheless, a legacy of agricultural pollutants persists in the A2+R2 connection, evidenced by *Rheinheimera* (p=0.004) and *Dechloromonas* (p=0.022), the latter known for degrading aromatic compounds. The sediment analysis reveals greater biogeochemical inertia (184 indicators) and incomplete substrate recovery. The altered sites (A) maintain a “dry land” edaphic footprint massively dominated by terrestrial actinobacteria such as *Nocardia*, *Streptosporangium*, *Pseudonocardia*, and *Actinophytocola*. Furthermore, the presence of *Nitrosospira* (stat=1.0, p=0.021) and archaea such as *Ca. Nitrososphaera* suggests active nitrification driven by residual fertilizer loads. Unlike the water, the restored sites (R) exhibit a hybrid state: although they begin to develop mature sediment characteristics shared with the well-preserved (e.g., *Woeseia* and *Gemmata* in WP+R), they retain a strong structural connection with the altered sites (Group A2+R2), marked by spore formers typical of soils such as *Bacillus*, *Paenibacillus*, and *Solibacillus*. This confirms that hydrological restoration has not yet erased the physical memory of the pasture soil, which coexists with natural functions such as the methane cycle (*Methylocaldum*) and cyanobacteria (*Cuspidothrix*) present in the sediments of well-preserved site.

#### Dutch Delta

In the Southwest Dutch Delta, the water analysis (265 indicators, p<0.05) revealed strong differentiation based on restoration maturity and hydrodynamics. The altered sites (A1+A2) reflect stagnation caused by physical barriers, with indicators such as *Roseospira* (stat=0.88, p=0.021), a photosynthetic bacterium, alongside *Cetobacterium* (stat=0.99, p=0.011) and *Defluviimonas* (p=0.008), fermenters associated with organic accumulation. Long-term restoration success is manifested in the massive convergence of the P2+R2 group (51 shared species), where genera such as *Rhizobacter* (p=0.003), *Silanimonas* (p=0.001), *Chryseobacterium* (p=0.003), and *Gemmatimonas* (p=0.002) validate that after 30 years the community is indistinguishable from the well-preserved. In contrast, recent restoration (R1, ∼4 years) shows only an incipient connection with its well-preserved (Group WP1+R1), limited to sulfur cycle bacteria such as *Dethiosulfatibacter* (p=0.001) and *Neptuniibacter* (p=0.001). Functionally, R1 sites are dominated by active redox cycles of post-inundation iron and sulfur release (*Ferrimonas*, *Desulfogranum*), whereas R2 exhibits greater degradative complexity with *Janthinobacterium* and *Sphingopyxis* (p=0.013). Finally, the well-preserved sites (WP) stand out for high natural primary productivity, marked by filamentous cyanobacteria such as *Pseudanabaena* and *Planktothrix* (p=0.011). The sediment analysis (187 indicators) confirms that sediment recovery is a slow process dependent on time and connectivity. The altered sites (A) present an impoverished community with few exclusive indicators, highlighting the cyanobacterium *Pleurocapsa* (p=0.037) in response to reduced resuspension. The recent restoration sites (R1) are in a “biogeochemical shock,” dominated by indicators of intense anoxia such as *Desulfosporosinus* (stat=0.91, p=0.002) and *Desulfobacula* (p=0.038), and halophilic fermenters like *Clostridiisalibacter* (stat=0.81, p=0.001), reflecting the rapid mineralization of terrestrial organic matter. Conversely, Long-term Restoration (R2) has allowed the establishment of stable and oxidative redox gradients, with the presence of *Gallionella* (p=0.006), *Sulfuricella* (p=0.006), and *Rhizobacter* (p=0.001). Long-term functional convergence is evidenced in the P2+R2 Group, which shares key species like *Desulfobulbus*, *Pseudazoarcus* (p=0.002), and *Gemmatimonas*, demonstrating the recovery of the fine biogeochemical structure. Additionally, a basal “delta microbiome” exists (Group A2+P2+R1+R2), composed of generalists such as *Sphingomonas*, *Hyphomicrobium*, and *Thiobacillus*, which transversely colonize sediments with hydrological connectivity.

#### Ria de Aveiro

In the Ria de Aveiro, water IndVal analysis (113 indicators, p<0.05) reflects the complex interaction between bioturbation caused by bait digging and estuarine hydrodynamics. Clustering patterns reveal a strong spatial dependence, where groups associated with altered sites (A), such as A1+P1, show clear indicators of sediment resuspension and deep anaerobic processes. Notably, the presence of *Methanococcoides* (p=0.047), a methanogenic archaeon whose detection in the water suggests that excavation releases methane and subsurface bacteria, stands out alongside *Agromyces* (p=0.047) and *Paenisporosarcina* (p=0.047), indicators of terrestrial input consistent with margin erosion. Restoration has created stable niches sharing characteristics with A (Group A1+P1 and A1+R1), including *Thiohalocapsa* (p=0.019), indicative of near-surface sulfurous conditions, and *Gramella* (p=0.019), associated with algal biomass degradation. However, promising signs of functional recovery emerge in the restored with well-preserved connection (A1+R1 and WP1), marked by *Castellaniella* (p=0.002), a denitrifier suggesting the re-establishment of key nitrogen cycle functions. Finally, the well-preserved sites (WP) exhibit halophilic stability with *Saccharospirillum* (p=0.016) and a complex trophic web evidenced by *Ca. Amoebophilus* (p=0.032). The sediment analysis (44 species by IndVal.g) shows critical specialization to sediment conditions driven by vegetation restoration (*Zostera*). The Restored Sites (R1) present the most distinctive “rhizosphere effect” signal, with the exclusive appearance of nitrifiers such as *Nitrosomonas* (p=0.009) and *Ammoniphilus* (p=0.007). This confirms that the roots of transplanted plants are oxygenating the sediment, creating niches for aerobic bacteria and root exudate degraders like *Flavobacterium* (p=0.038) and *Pseudomonas* (p=0.024). In contrast, the altered sites (A) reflect the impact of continuous mechanical disturbance, dominated by strict anaerobes such as *Aerophobus* (p=0.016), *Sumerlaea* (p=0.004), and *Vallitalea* (p=0.016) in A2, and an unstable sulfur cycle indicated by *Desulfurivibrio* (p=0.007) in A1. Despite these differences, a functional estuarine “core” (Group A+P+R) exists, shared by all sites and composed of *Limnobacter*, *Sedimenticola*, *Desulfococcus*, and *Ca. Nitrosoarchaeum*, maintaining basal sulfur and nitrogen cycles. The ultimate success of restoration is validated in the P+R connection, where the shared presence of *Ca. Tenderia* (p=0.014), a key denitrifier, and *Algoriphagus* (p=0.032), indicates that restored sites are recovering the nitrogen metabolism capacity.

## Discussion

Our results provide a partial validation of the initial hypothesis, confirming that ecological restoration effectively reconfigures the microbial community, however this process is not synchronous between water and sediment compartments. Differences in environmental variables and ecosystem types reveal a biogeographical pattern in the obtained results and constitutes the primary hierarchical filter structuring microbial diversity at the continental scale. The sharp segregation of communities in ordination analyses and the formation of site-exclusive clusters in heatmaps, is evidence that environmental management exerts a secondary yet functionally critical selective pressure that reveals a fundamental decoupling in ecosystem recovery dynamics (Martiny et al., 2006; Hanson et al., 2012; Miralles et al., 2025). The data indicate that ecological restoration does not affect compartments simultaneously; while the bacterioplankton of in water exhibits high plasticity and rapid response capacity, frequently converging towards reference states (well-preserved) following the restitution of hydrological or salinity conditions, as observed in the recovery of halophiles in Marja del Moros (VA) or the elimination of eutrophication indicators in the Danube Delta, the sediment manifests a marked “ecological memory” or hysteresis (Moreno-Mateos et al., 2012; Suding et al., 2004). This sediment inertia is characterized by the persistence of microbial guilds inherited from the alteration period (e.g., shared A2+R2 clusters in the Danube and Dutch Delta), suggesting that the biogeochemical legacy of prior land use creates a temporal barrier that delays the functional reconfiguration of the substrate far beyond the implementation of physical restoration measures (Allison & Martiny, 2008).

Results reveal that microbial community structure is primarily governed by environmental characteristics and ecosystem type, which define the “core” identity of each ecosystem over local conservation status (Lozupone & Knight, 2007). In water, there is a distinct segregation: Marjal dels Moros (VA) is isolated with a distinctly structured microbial community in the multivariate analysis, likely due to its hypersaline conditions and confined hydrological management, whereas systems with greater continental and freshwater influence, such as the Danube Delta and Curonian Lagoon, tend to cluster in multivariate space. Conversely, sediments delineate a continuous biogeographical gradient connecting Mediterranean systems (Marjal del Moros (VA), Camargue (CA)) to Atlantic ones (Ria de Aveiro (RI), Dutch Delta (DU)), broken only by the biogeochemical singularity of the Curonian Lagoon, which emerges as the most distinctive sediment. This pattern confirms that regional factors such as latitude, marine connectivity, and salinity regimes impose the basal assembly rules upon which management pressures subsequently act (Lindström & Langenheder, 2012).

However, despite this marked geographic differentiation, we observed a critical phenomenon of “functional convergence” or biotic homogenization in altered sites (McKinney & Lockwood, 1999). Regardless of whether the wetland is in the Mediterranean, the Atlantic, or the Black Sea, degradation appears to homogenize certain functional traits of the microbiota, favoring the proliferation of strict anaerobic guilds and bacteria associated with dysfunctional sulfur cycles. This is evidenced by the recurrent appearance of indicator genera such as *Desulfomonile* and fermenters like *Clostridium* or *Propionivibrio* in the altered sediments of sites as distant as VA and CA. This ubiquity of anoxia and organic load markers suggests that hydrological alteration and eutrophication force microbial communities originally distinct towards a common metabolic state dominated by energetic inefficiency and the production of toxic reduced compounds (Lamers et al., 2013). This validates the hypothesis that, while healthy ecosystems are unique in their diversity (as seen in well-preserved sites), altered ecosystems tend to functionally resemble one another.

The observed temporal decoupling manifests as a phenomenon of ecological hysteresis, where the recovery trajectory of the microbial community does not mirror its degradation path, particularly in the sediment (Beisner et al., 2003). While water responds almost immediately to the restoration of physical drivers, evidenced by the rapid convergence of pelagic communities in restored and well-preserved sites in Marjal del Moros (VA) and the Danube Delta (DA), sediments act as reservoirs of “ecological memory.” This inertia is particularly visible in the Danube Delta, where, despite hydrological reconnection, sediments in restored sites continue to harbor a significant core of terrestrial actinobacteria and spore-formers like *Bacillus*, remnants of prior agricultural and livestock use. Similarly, in Marjal dels Moros (VA) although the water has been restored towards saline conditions, the sediment in the restored subsite maintains a robust taxonomic connection with altered sites (Group A2+R1), indicating that refractory organic matter accumulated in sediment during decades of eutrophication continues to condition food webs long after water quality has improved. This finding underscores that hydrological restoration is a necessary but insufficient condition to erase the soil’s biogeochemical legacy, which requires decadal timescales for complete reconfiguration (Moreno-Mateos et al., 2015).

Regarding the functional analysis via bioindicators, and more specifically for carbon dynamics and GHG metabolisms, indicator species analysis reveals that hydrological alteration transforms wetlands into potential active CH_4_ emitters, a function that restoration seeks to reverse (Bridgham et al., 2013). In the altered sediments of the Danube Delta and the tidally renewed water of the Ria de Aveiro, the significant detection of strict methanogens such as *Methanothrix* and *Methanococcoides*, respectively, evidence active methane production driven by anoxia and the availability of fermentable substrates. Conversely, the presence of methanotrophic bacteria such as *Methylophaga* in the well-preserved sites of the Camargue and Dutch Delta suggests that functional ecosystems develop an efficient “biofilter” capacity to consume generated CH_4_ (Dean et al., 2018). However, the persistence of methanotrophs is indicative of high methane loading, such as *Methyloparacoccus* in the altered waters of the Curonian Lagoon, and warns on how eutrophication can exacerbate these emissions.

The composition of sulfur cycle bacteria acts as an accurate thermometer of the redox state of the sediment (Lamers et al., 2013). In the altered sites of Marjal del Moros and the Camargue, the dominance of sulfate reducers such as *Desulfomonile* and *Desulfovibrio* indicate strict anoxic conditions and potential accumulation of hydrogen sulfide. This prevalence of sulfate-reducing bacteria (SRB) has profound implications for the ecosystem’s GHG budget. Since sulfate reduction is thermodynamically more favorable than methanogenesis, these bacteria effectively outcompete methanogenic archaea for common electron donors (e.g., acetate and hydrogen), thereby acting as a biotic mechanism that suppresses potential methane (CH_4_) emissions (Lovley & Klug, 1983). However, this competitive suppression presents a critical ecological trade-off: while it may limit the release of CH_4_, it drives the accumulation of phytotoxic sulfides which can inhibit vegetation recovery, illustrating the complex functional constraints inherent to altered biogeochemical states (Koch et al., 1990). Successful ecological restoration is manifested in the appearance of phototrophic and chemolithotrophic sulfur-oxidizing bacteria, such as *Chromatium* and *Thiovirga*, exclusive to restored sites (R2) in the Camargue. This colonization signals the re-establishment of functional redox gradients where sulfide (energy source) and oxygen or light coexists, allowing for sediment support of greater biodiversity.

Regarding water quality Indicators, bacterial community structure sharply reflects the trophic state of the system, differentiating between water dominated by detritus recycling and those sustained by primary production. In the Danube Delta, altered sites are marked by indicators of organic and fecal contamination like *Shewanella* and *Arcobacter*, denoting a heterotrophic system saturated with labile organic matter. In contrast, hydrological restoration in this same delta has fostered a shift towards autotrophy, evidenced by the proliferation of picocyanobacteria such as *Cyanobium* and *Synechocystis* in restored site (R1), typical of clearer waters with functional food chains. This pattern of microbiological “cleansing” validates the effectiveness of restoration measures in reversing states of severe eutrophication inherited from intensive agriculture.

The case of the Dutch Delta critically illustrates how restoration maturity determines microbiome functionality in sediment. Recently restored site (R1, ∼4 years) are dominated by genera such as *Ferrimonas* and *Desulfosporosinus*, indicators of intense iron and sulfate reduction processes characteristic of newly flooded terrestrial soils undergoing a reductive “shock.” In contrast, older restoration (R2, >30 years) has allowed for the development of a complex community with species like *Desulfobulbus* and *Janthinobacterium*, indistinguishable from reference sites. This confirms that the stabilization of biogeochemical cycles in sediment is a process operating on decadal scales, warning that the evaluation of restoration success must not be premature.

The implications of these findings for wetland management and climate mitigation are profound, validating specific restoration strategies while issuing a critical warning regarding the limitations of conventional monitoring protocols. Our data confirm the efficacy of targeted interventions, such as salinity management in Marjal dels Moros (VA), which has proven to be a potent tool for “resetting” the pelagic microbial community, eliminating pathogens associated with freshwater eutrophication and favoring the recruitment of functional halophilic taxa, or active revegetation in the Ria de Aveiro, where plant reintroduction (*Zostera*) has been determinant in oxygenating the sediment and reactivating nitrification (e.g., *Nitrosomonas*) via the “rhizosphere effect” (Brodersen et al., 2015; Olsen et al., 2016). However, the observed dissociation between biological matrices warns us that assessing restoration success based solely on water quality may lead to diagnoses of “false success”; systems with apparently recovered water, as in the Danube Delta, may conceal sediments that continue to function as active methane reactors and reservoirs of inherited nutrients due to their biogeochemical inertia. Consequently, it is imperative that future management programs incorporate molecular analysis of sediments as an indispensable success indicator, thereby ensuring that functional recovery encompasses not only surface aesthetics or water chemistry, but also the deep processes in sediment that regulate the ecosystem’s true long-term climate resilience.

The application of molecular tools has allowed us to go beyond mere structural assessments of restoration, although this study does not examine temporal dynamics at a single study site. The experimental design comparing old, well-preserved, and restored areas allows us to establish that the recovery of coastal wetlands after restoration processes must consider that water and sediment behave very differently in this restoration process. The rapid rehabilitation of water contrasts with the tenacious biological inertia in the sediment. Our results demonstrate that while it is possible to re-establish hydrological conditions favoring healthy pelagic communities in the short term, sediments act as archives of historical degradation, retaining dysfunctional microbial signatures that may take decades to dissipate. Therefore, ecological restoration must be understood not only as the reconstruction of visible landscapes but as the patient re-engineering of invisible biogeochemical functions, recognizing that the ecosystem’s resilience will depend on our capacity to sustain interventions long enough for the sediment to finally overcome its memory of alteration.

## Conclusions

After ecological restoration, the diversity and complexity of the microbial community partially recover towards natural reference standards, although the restored sites often retain distinct signatures that differ from both well-preserved and altered, suggesting a transitional recovery trajectory rather than an immediate return to the characteristic community of well-preserved sites.

Compared to altered and well-preserved conditions, the impact of restoration on prokaryotic community structure exhibits marked differences between water and sediment: while bacterioplankton in the water shows high resilience and rapid convergence with natural sites, the sediment microbiome displays significant “ecological memory,” maintaining an altered community structure due to the inertia in sediment. The restoration trajectories can be followed by bacterial genera that act as indicators for specific functional states, allowing for the molecular diagnosis of ecosystem recovery serving as sentinels to validate the re-establishment of biogeochemical cycles beyond mere physicochemical parameters.

Restoration promotes structural shifts with functional implications on the metabolic pathways associated with greenhouse gas emissions, specifically by modulating the balance between syntrophic consortia, methanogens, and methanotrophs, which is critical for shifting the ecosystem function from a GHG source back to a GHG sink.

An integrated approach is essential for coastal wetland restoration, ensuring that interventions restore critical biogeochemical processes and support the assembly of prokaryotic communities in water and sediments across European coastal wetlands, thereby enhancing resilience under climate change.

## AuthorContributions

**Picazo, A.**: Writing-Original Draft, Conceptualization, Data curation, Formal analysis, Investigation, Methodology, Resources. **Rochera, C.**: Conceptualization, Data curation, Investigation, Methodology, Resources. **Morant, D.**: Conceptualization, Data curation, Methodology, Resources. **Pedrón, M.**: Investigation. **Adamescu, M.**: Investigation, Resources. **Ambrosio, R.**: Investigation, Project administration. **Attermeyer, K.**: Methodology, Resources. **Santinelli, C.**: Data curation, Formal analysis, Investigation, Resources, Validation. **Tropea, C.**: Data curation, Formal analysis, Investigation, Resources. **Bachi, G.**: Investigation, Methodology. **Bègue, N.**: Investigation. **Bučas, M.**: Data curation, Formal analysis, Investigation, Resources. **Cabrera-Brufau, M.**: Data curation, Resources. **Camacho Santamans, A.**: Resources. **Carballeira, R.**: Investigation. **Carloni, M.**: Formal analysis, Investigation. **Cavalcante, L.**: Data curation, Resources. **Cazacu, C.**: Methodology, Resources. **Checcucci, G.**: Investigation, Methodology. **Coelho, J.P.**: Data curation, Methodology, Resources. **Dinu, V.**: Investigation, Resources. **Evangelista, V.**: Formal analysis, Investigation. **Gintauskas, J.**: Investigation. **Giuca, R.**: Investigation, Resources. **Guelmami, A.**: Investigation, Project administration, Resources. **Guerrazzi, M.**: Formal analysis, Investigation. **Hilaire, S.**: Investigation. **Kataržytė, M.**: Investigation. **Lillebø, A.I.**: Conceptualization, Data curation, Methodology, Project administration, Resources. **Lizán, R.**: Formal analysis, Investigation. **Minaudo, C.**: Conceptualization, Data curation, Methodology, Resources. **Misteli, B.**: Conceptualization, Data curation, Methodology, Resources. **Montes-Pérez, J.**: Resources. **Obrador, B.**: Conceptualization, Methodology, Resources. **Oliveira, B.R.F.**: Data curation, Methodology, Resources. **Oliveira, V.H.**: Resources. **Petkuvienė, J.**: Data curation, Formal analysis, Investigation, Resources. **Racoviceanu, T.**: Investigation, Resources. **Ribeiro, S.**: Data curation, Methodology, Resources. **Ronse, M.**: Investigation, Resources. **Sousa, A.**: Data curation, Methodology, Resources. **Suykerbuyk, W.**: Formal analysis, Investigation. **Tiškus, E.**: Investigation. **Vaičiūtė, D.**: Data curation, Formal analysis, Investigation, Resources. **Valsecchi, S.**: Formal analysis, Investigation. **van Puijenbroek, M.**: Data curation, Investigation, Resources. **Andreu, O.**: Formal analysis. **Yegon, M.**: Resources. **von Schiller, D.**: Conceptualization, Methodology, Resources. **Camacho, A.**: Conceptualization, Data curation, Formal analysis, Funding acquisition, Investigation, Methodology, Project administration, Supervision, Validation, Writing - review & editing.

## Funding

This research has received funding from the project RESTORE4Cs - Modelling RESTORation of wEtlands for Carbon pathways, Climate Change mitigation and adaptation, ecosystem services, and biodiversity, Co-benefits (DOI: 10.3030/101056782), co-funded by the European Union under the Horizon Europe research and innovation programme (Grant Agreement ID: 101056782). The views and opinions expressed are those of the author(s) only and do not necessarily reflect those of the European Union or the granting authority. Neither the European Union nor the granting authority can be held responsible for them. Work by the University of Valencia was also co-supported by projects CLIMAWET-CONS (PID2019-104742RB-I00), funded by the Agencia Estatal de Investigation of the Spanish Government, and by the project ECCAEL (PROMETEO CIPROM-2023-031), funded by the Generalitat Valenciana, both granted to Antonio Camacho.

## Acknowledgements

UAveiro team acknowledge the support of FCT – Fundação para a Ciência e a Tecnologia I.P., to CESAM-Centro de Estudos do Ambiente e do Mar, funded by national funds through FC, under the project references UID/50017/2025 (doi.org/10.54499/UID/50017/2025) and LA/P/0094/2020 (doi.org/10.54499/LA/P/0094/2020). Work by the University of Valencia was also co-supported by projects CLIMAWET-CONS (PID2019-104742RB-I00), funded by the Agencia Estatal de Investigation of the Spanish Government, and by the project ECCAEL (PROMETEO CIPROM-2023-031), funded by the Generalitat Valenciana, both granted to Antonio Camacho.

## Conflict of interest statement

The authors declare that they have no conflict of interest

## Data availability

All sequence data from this study have been deposited in the Sequence Read Archive (SRA) of the National Center for Biotechnology Information (NCBI), BioProject accession number SUB15896998.

## Declaration of generative AI and AI-assisted technologies in the manuscript preparation process

During the preparation of this work the authors used ChatGPT-5.2 to identify potential improvements in text readability and for coding syntax support during data processing. After using this tool/service, the authors reviewed and edited the content as needed and took full responsibility for the content of the published article.

## Notes

### Competing Interest Statement

The authors have declared no competing interest.

